# Multiple evidences suggest *sox2* as the main driver of a young and complex sex determining ZW/ZZ system in turbot (*Scophthalmus maximus*)

**DOI:** 10.1101/834556

**Authors:** Paulino Martínez, Diego Robledo, Xoana Taboada, Andrés Blanco, Antonio Gómez-Tato, Blanca Álvarez-Blázquez, Santiago Cabaleiro, Francesc Piferrer, Carmen Bouza, Ana M. Viñas

**Affiliations:** University of Santiago de Compostela; University of Edinburgh; Spanish Institute of Oceanography (IEO); Aquaculture Cluster; Institute of Marine Sciences

**Keywords:** Turbot, sex determination evolution, GWAS, sox2, sexual differentiation, lncRNA

## Abstract

A major challenge in evolutionary biology is to find an explanation for the variation in sex-determining (SD) systems across taxa and to understand the mechanisms driving sex chromosome differentiation. We studied the turbot, holding a ZW/ZZ SD system and no sex chromosome heteromorphism, by combining classical genetics and genomics approaches to disentangle the genetic architecture of this trait. RAD-Seq was used to genotype 18,214 SNPs on 1,135 fish from 36 families and a genome wide association study (GWAS) identified a ∼ 6 Mb region on LG5 associated with sex (P < 0.05). The most significant associated markers were located close to *sox2*, *dnajc19* and *fxr1* genes. A segregation analysis enabled narrowing down the associated region and evidenced recombination suppression in a region overlapping the candidate genes. A Nanopore/Illumina assembly of the SD region using ZZ and WW individuals identified a single SNP fully associated with Z and W chromosomes. RNA-seq from 5-90 day-old fish detected the expression along the gonad differentiation period of a short non-coding splicing variant (ncRNA) included in a vertebrate-conserved long non-coding RNA overlapping *sox2*. qPCR showed that *sox2* was the only differentially expressed gene between males and females at 50-55 days post fertilization, just prior the beginning of gonad differentiation. More refined information on the involvement of secondary genetic and environmental factors and their interactions on SD was gathered after the analysis of a broad sample of families. Our results confirm the complex nature of SD in turbot and support *sox2* as its main driver.

## INTRODUCTION

Sex is one of the most intriguing topics from evolutionary and development perspectives. Sex determination (SD), the process that establishes the sex of an organism at the initial stages of development, affects sex ratio and hence, it has profound implications on demography of populations. Gonad differentiation (GD), the pathway by which an undifferentiated primordium develops into an ovary or testis, is also essential to understand how sex is controlled and shifts from one system to another along evolution.

Theories on the evolution and architecture of SD have been largely influenced by studies on mammals, birds and *Drosophila*, characterized by a highly conserved master SD gene (SDg). However, data from ectothermic vertebrates have demonstrated a sharply different picture (Devlin and Nagahama, 2002; Martínez et al., 2014; Capel et al., 2017). Fish show highly diverse SD systems, including analogous models to the XY of mammals and ZW of birds, but also more complex mechanisms involving multiple sex chromosomes. Regardless of their SD system, chromosome heteromorphisms are rare in this vertebrate group (Cioffi et al., 2017). Underlying this diversity, an important number of different SDg have been reported, both involving classical transcription factors, such as *dmy* (Y-specific DM-domain) (Matsuda et al., 2002) or *sox3* (SRY-related HMG-box) (Takehana et al., 2014), as well as transforming growth factor-related genes, such as *gsdf* (gonadal soma-derived growth factor on the Y chromosome) (Myosho et al., 2012) or *amh* (anti-Mullerian hormone) (Hattori et al., 2012; Pan et al., 2019) and its receptor *amhr2* (Kamiya et al., 2012), but also other unexpected genes, such the interferon-related *sdY* (Yano et al., 2013), and more recently, *bcar1* (Bao et al., 2019) and *hsd17b1* (Koyama et al., 2019), related to the steroidogenic pathway. Furthermore, quantitative trait loci (QTL) studies in other fish suggest that other non-orthologous genomic regions could be involved in SD, the current list being likely expanded in the next years (Peichel et al., 2004; Ser et al., 2010; Palaiokostas et al., 2013; Star et al., 2016). In contrast with the highly canalized systems of homeothermic vertebrates, involving a strictly hierarchical and highly conserved developmental pathway, the multiple existing options to recruit new SD genes shown by fish suggest a more relaxed hierarchy, and even a more flexible downstream pathway until the gonad fate is established (Martínez et al., 2014).

The enormous SD diversity in fish is associated with a remarkable SD evolutionary turnover and, in fact, closely related species have demonstrated different SD systems, such as the case of the genera *Oryzias* (Matsuda and Sakaizumi, 2016) and *Oreochromis* (Baroiller and D’Cotta, 2019). Nonetheless, a remarkable conservation of SD has also been reported in other groups, such as the genus *Seriola* (Koyama et al., 2019) or the order Salmoniformes (Yano et al., 2016). It has been hypothesized that intraspecific variability might facilitate the high SD evolutionary turnover of fish (Martínez et al., 2014) and accordingly, these authors suggested focusing not only on the master SDg, but also on the genetic variation within species to obtain a more comprehensive picture of the SD genetic architecture. In fact, the limited number of studies following this approach have disclosed significant genetic variation within species, even in those where a major SDg has been reported (Hermida et al., 2013; Lozano et al., 2013; Parnell and Streelman, 2013).

The environment adds another layer of complexity, since fish SD can also be influenced by a range of environmental factors. Accordingly, fish SD has been described as a continuum from strict environmental SD (ESD) to pure genetic SD (GSD), involving minor and major genetic factors (Penman and Piferrer, 2008). It is difficult to encompass the diversity of fish SD within a conceptual genetic framework, but probably the closest would be a complex threshold trait and consequently, quantitative genetic tools would be the most appropriate for its dissection (Martínez et al., 2014; Capel, 2017).

The turbot (*Scophthalmus maximus*) is an important farmed fish in Europe (∼10,000 tons) and China (∼60,000 tons, Robledo et al., 2018). This species shows one of the highest growth rate sex dimorphisms among marine fish in favor of females (Piferrer et al., 2004), hence, the interest of turbot farmers to produce all female stocks. Since sex cannot be identified until maturation, molecular tools have been sought for precocious sexing of farmed turbot (Martínez et al., 2014). Turbot shows a SD system compatible with ZW/ZZ (Haffray et al., 2009; Martínez et al., 2009), but no sex chromosome heteromorphism has been detected (Bouza et al., 1994; Cuñado et al., 2002). Previous data suggest that the differential SD region in this species is small and that its SD mechanism is recent in evolutionary terms (Taboada et al., 2014). A major SD region on LG5 and three additional suggestive sex-related QTL (LG6, LG8 and LG21) were identified through a genome scan using five families (Martínez et al., 2009; Hermida et al., 2013) and a medium density genetic map (Bouza et al., 2007). Furthermore, temperature influence has also been demonstrated in turbot (Haffray et al., 2009; Robledo et al., 2015), so its SD mechanism approximates to a complex trait. A microsatellite sex associated marker (SmaUSC-E30) found in the proximal region of LG5 allowed accurate sexing of 94.7% offspring (Martínez et al., 2009) and it has been routinely used by the industry to obtain all-female populations (Martínez et al., 2014). Mapping of sex-related genes and mining around the major SD region using the turbot genome (Figueras et al., 2016) led to the identification of candidate genes at the major (LG5: *sox2 dnajc19* and *fxr1*; Taboada et al., 2014) and minor (LG6: *cyp19b*; LG21: *sox9* and *sox17*; Viñas et al., 2012) QTL, but none of them seemed to be the SDg of the species (Taboada et al., 2014). The first evidences of gonad development in turbot were detected at 65 days post fertilization (dpf), where *gsdf1* in both sexes and *vasa*, especially in females, were up-regulated. By 90 dpf, the gonad seems to be transcriptionally differentiated, and *cyp19a1a* and *amh* show differential expression in females and males, respectively (Robledo et al., 2015; Ribas et al., 2016).

Here, we combined functional and genome wide association studies (GWAS), as well as classical segregation and recombination analyses in a large sample of ∼1200 turbot from 36 families with the aim of: i) identifying the major SDg and eventually the causal mutation differentiating males and females; ii) characterizing the ZW differential region and its genetic properties within an evolutionary framework; iii) refining gonad development information establishing the critical window where gonad fate is established; and iv) assessing interfamily variation underlying SD. We identify sox2 as the most likely SD candidate in turbot, proposed suggestive hypotheses about how this gene is differentially regulated in males and females, and finally, explore in more detail how other minor QTLs and environmental signals underlie SD in a species where the Z and W chromosomes hardly differ.

## RESULTS

### Identifying the SD region and evaluating its influence on phenotypic sex

The study population was composed by 1135 fish distributed across 36 families (from 23 to 38 fish per family) founded from 23 dams and 23 sires, hence including several half-sib families (HS) sharing either the mother or the father (Table 1). The sex ratio in the whole population adjusted to 1:1 (563 females and 592 males; P = 0.394); most families showed balanced sex ratios, excluding one male-biased (F53) and two female-biased (F12 and F33) (P < 0.00005; Figure S1). All animals were genotyped using 2b-RAD for 18,214 SNPs.

**Table 1:**
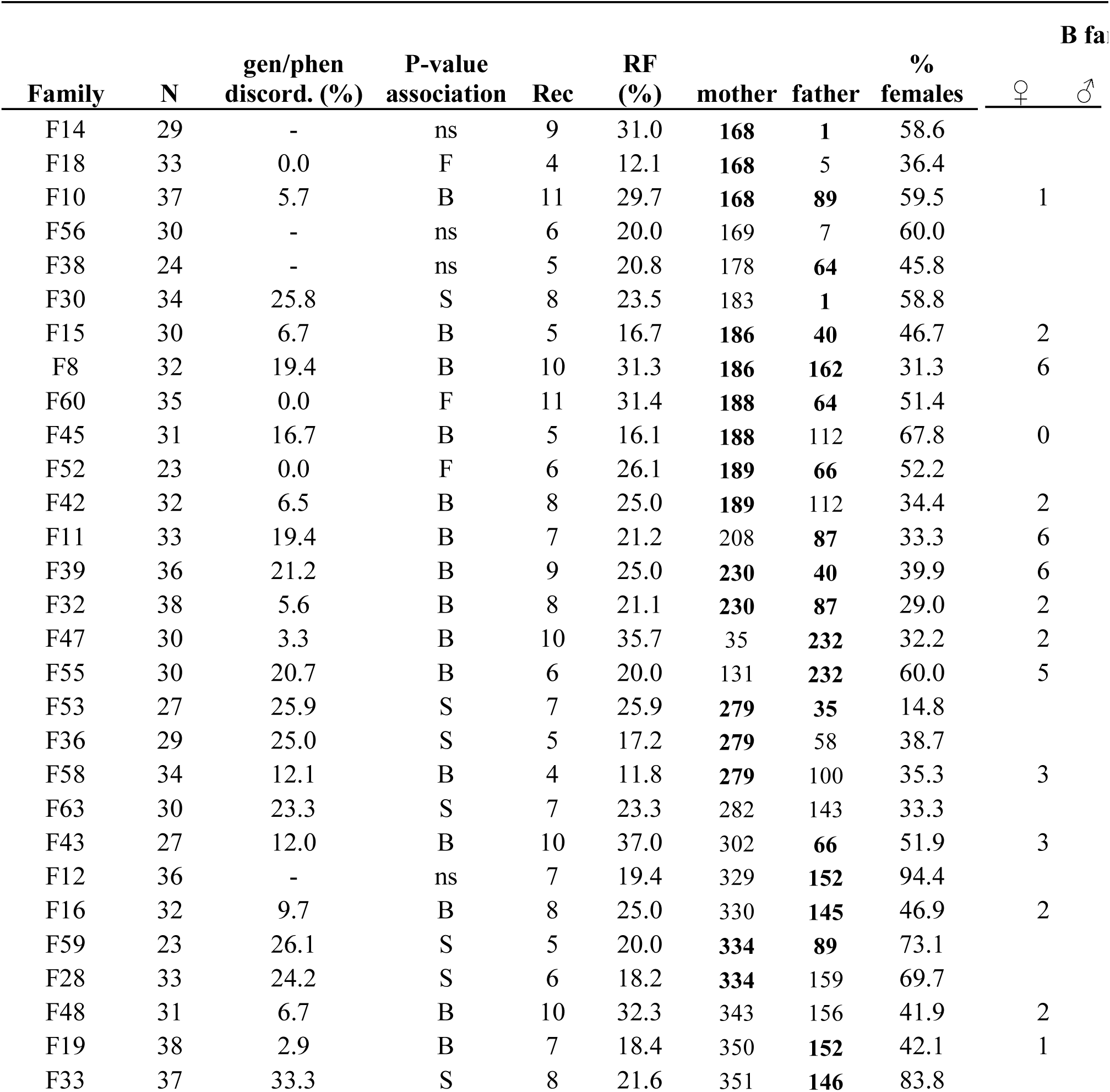
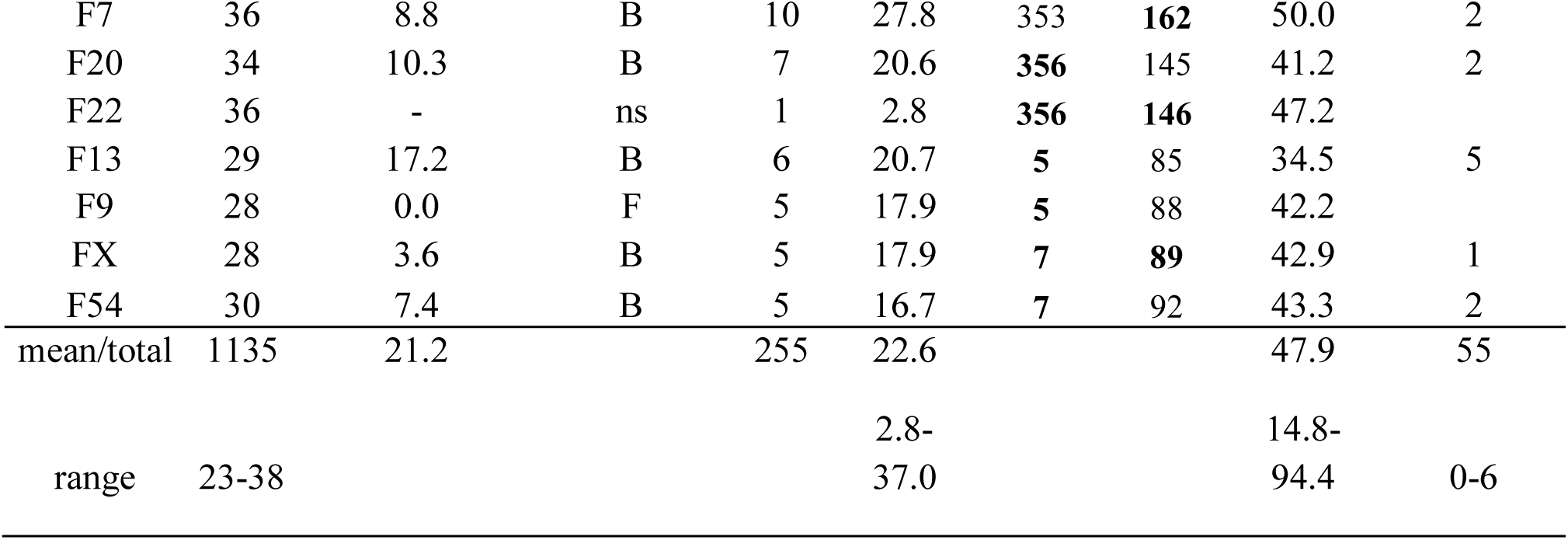
Segregation analysis at LG5 in 36 *S*. *maximus* families analyzed to look for association with of offspring per family; gen/phen discord.: % of discordances between genetic and phenotypic sex at phen sex association (F: full, no discordances between phenotypic and genetic sex; B: P < 0,05 after B P < 0,05; ns: not significant association); Rec: number of recombinants; RF: recombination frequency genetic females that are phenotypic males and ♂ → ♀ number of genetic males that are phenotyp bold are shared by different families (half-sibs).

| Family | N | gen/phen<br>discord. (%) | P-value<br>association | Rec | RF<br>(%) | mother | father | %<br>females | B families |  |
| --- | --- | --- | --- | --- | --- | --- | --- | --- | --- | --- |
|  |  |  |  |  |  |  |  |  | ♀ | ♂ |
| F14 | 29 | - | ns | 9 | 31.0 | **168** | **1** | 58.6 |  |  |
| F18 | 33 | 0.0 | F | 4 | 12.1 | **168** | 5 | 36.4 |  |  |
| F10 | 37 | 5.7 | B | 11 | 29.7 | **168** | **89** | 59.5 | 1 |  |
| F56 | 30 | - | ns | 6 | 20.0 | 169 | 7 | 60.0 |  |  |
| F38 | 24 | - | ns | 5 | 20.8 | 178 | **64** | 45.8 |  |  |
| F30 | 34 | 25.8 | S | 8 | 23.5 | 183 | **1** | 58.8 |  |  |
| F15 | 30 | 6.7 | B | 5 | 16.7 | **186** | **40** | 46.7 | 2 |  |
| F8 | 32 | 19.4 | B | 10 | 31.3 | **186** | **162** | 31.3 | 6 |  |
| F60 | 35 | 0.0 | F | 11 | 31.4 | **188** | **64** | 51.4 |  |  |
| F45 | 31 | 16.7 | B | 5 | 16.1 | **188** | 112 | 67.8 | 0 |  |
| F52 | 23 | 0.0 | F | 6 | 26.1 | **189** | **66** | 52.2 |  |  |
| F42 | 32 | 6.5 | B | 8 | 25.0 | **189** | 112 | 34.4 | 2 |  |
| F11 | 33 | 19.4 | B | 7 | 21.2 | 208 | **87** | 33.3 | 6 |  |
| F39 | 36 | 21.2 | B | 9 | 25.0 | **230** | **40** | 39.9 | 6 |  |
| F32 | 38 | 5.6 | B | 8 | 21.1 | **230** | **87** | 29.0 | 2 |  |
| F47 | 30 | 3.3 | B | 10 | 35.7 | 35 | **232** | 32.2 | 2 |  |
| F55 | 30 | 20.7 | B | 6 | 20.0 | 131 | **232** | 60.0 | 5 |  |
| F53 | 27 | 25.9 | S | 7 | 25.9 | **279** | **35** | 14.8 |  |  |
| F36 | 29 | 25.0 | S | 5 | 17.2 | **279** | 58 | 38.7 |  |  |
| F58 | 34 | 12.1 | B | 4 | 11.8 | **279** | 100 | 35.3 | 3 |  |
| F63 | 30 | 23.3 | S | 7 | 23.3 | 282 | 143 | 33.3 |  |  |
| F43 | 27 | 12.0 | B | 10 | 37.0 | 302 | **66** | 51.9 | 3 |  |
| F12 | 36 | - | ns | 7 | 19.4 | 329 | **152** | 94.4 |  |  |
| F16 | 32 | 9.7 | B | 8 | 25.0 | 330 | **145** | 46.9 | 2 |  |
| F59 | 23 | 26.1 | S | 5 | 20.0 | **334** | **89** | 73.1 |  |  |
| F28 | 33 | 24.2 | S | 6 | 18.2 | **334** | 159 | 69.7 |  |  |
| F48 | 31 | 6.7 | B | 10 | 32.3 | 343 | 156 | 41.9 | 2 |  |
| F19 | 38 | 2.9 | B | 7 | 18.4 | 350 | **152** | 42.1 | 1 |  |
| F33 | 37 | 33.3 | S | 8 | 21.6 | 351 | **146** | 83.8 |  |  |

|  |  |  |  |  |  |  |  |  |  |
| --- | --- | --- | --- | --- | --- | --- | --- | --- | --- |
| F7 | 36 | 8.8 | B | 10 | 27.8 | 353 | **162** | 50.0 | 2 |
| F20 | 34 | 10.3 | B | 7 | 20.6 | **356** | 145 | 41.2 | 2 |
| F22 | 36 | - | ns | 1 | 2.8 | **356** | **146** | 47.2 |  |
| F13 | 29 | 17.2 | B | 6 | 20.7 | **5** | 85 | 34.5 | 5 |
| F9 | 28 | 0.0 | F | 5 | 17.9 | **5** | 88 | 42.2 |  |
| FX | 28 | 3.6 | B | 5 | 17.9 | **7** | **89** | 42.9 | 1 |
| F54 | 30 | 7.4 | B | 5 | 16.7 | **7** | 92 | 43.3 | 2 |
| mean/total | 1135 | 21.2 |  | 255 | 22.6 |  |  | 47.9 | 55 |
| range | 23-38 |  |  |  | 2.8-37.0 |  |  | 14.8-94.4 | 0-6 |

A genome-wide association study (GWAS) using the whole population data was performed to analyze the genetic architecture of turbot SD. A major QTL was detected at LG5 with 80 significant markers at genome-wide level (Figure 1), while weak signals were detected at other LGs, specifically at LG22, where one marker was significant at chromosome-wide level (P < 0.00002; Table S1). This picture is essentially concordant with previous reports (Martínez et al., 2009; Hermida et al., 2013). The genome-wide significant markers at LG5 spanned over a region of ∼7.5 Mb and over five scaffolds of the turbot genome (Figueras et al., 2016), with the two most significant SNP markers placed 420 Kb away (Figure 1; Table S1). These two markers were in the vicinity of *sox2*, *dnajc19* and *fxr1*, the three candidate genes at the SD region reported by Taboada et al. (2014; Figure 1), encoding for a DNA-binding transcription factor, a heat shock protein and a RNA-binding protein, respectively. The highest genetic differentiation coefficient (F_ST_) between male and female subpopulations was 0.193 at SNP-129698, located within an intron of *dnajc19*, and close to SmaUSC-E30, the marker used for sexing in turbot farms (Martínez et al., 2014). It should be noted that the maximum F_ST_ between males (ZZ) and females (ZW) for a diagnostic locus of the W and Z chromosomes would be 0.5 (Utsunomía et al., 2017).

**Figure 1:**
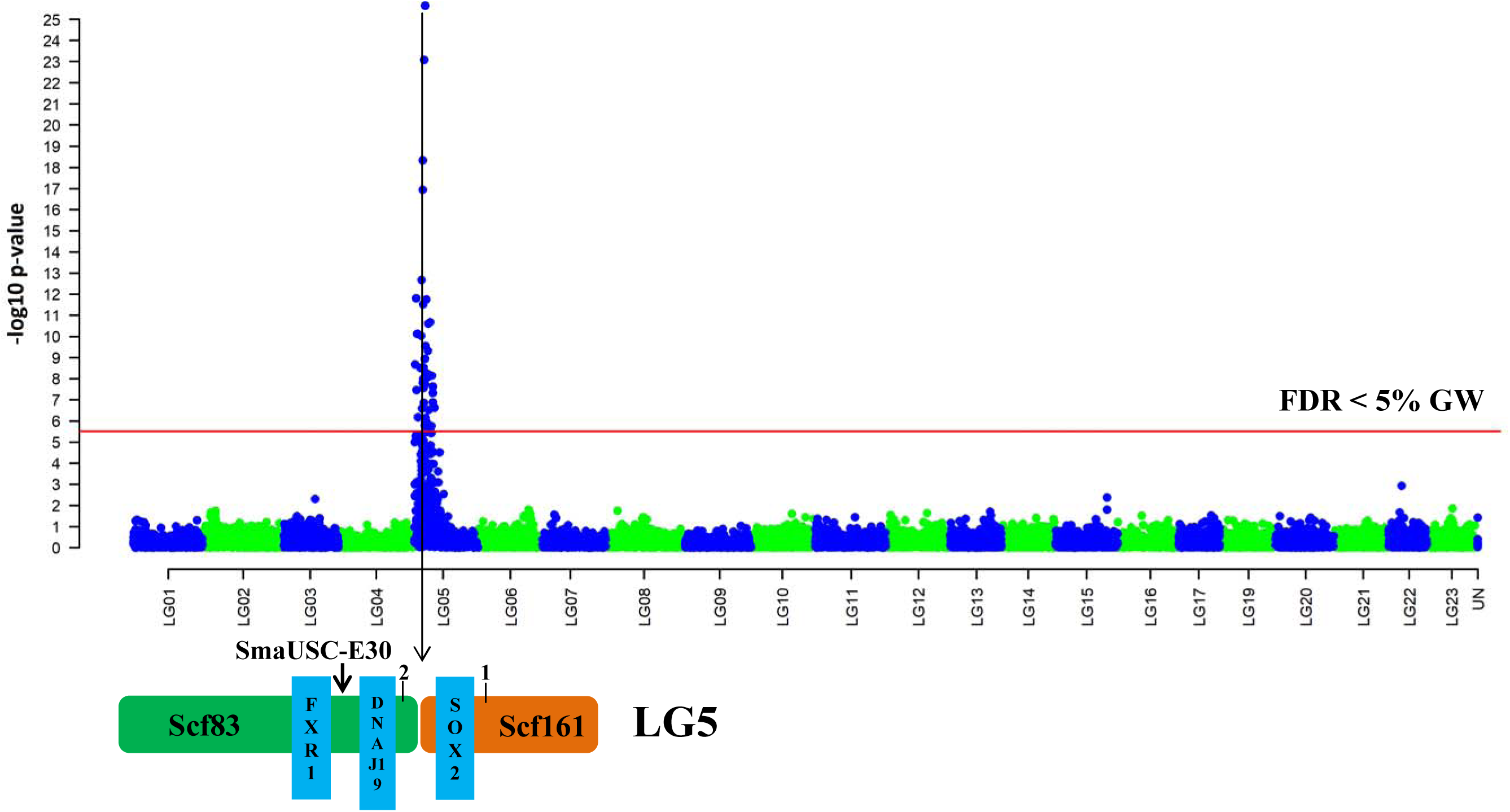
Genome-wide association study (GWAS) on turbot (*Scophthalmus maximus*) sex using the whole population data. Below the Manhattan plot is shown a zoom on the main associated genomic region at LG5 including the sex-associated microsatelite and the candidate genes previously reported by Martínez et al. (2009) and Taboada et al. (2014), respectively. Numbers 1 and 2 represent the first two highest significant sex-associated SNPs detected in the GWAS.

GWAS was also performed within each family, and markers at LG5 were associated with sex at genome- or chromosome-wide level in most families (Figure 2; Table S2). However, in some families no evidence of association with LG5 was detected and the highest associated marker pertained to other LGs (Table S2), although none of them significant at genome- or chromosome-wide level. Two families showed suggestive signals of association with LG22 (F19 and F36), which reinforces the observation with the whole population data, and other three with LG1 (F7), LG15 (F48) and LG20 (F43). It should be noted that alternative associations were not a matter of a single marker, but of several SNPs with p-values above the background denoting a trend for the region. The sample size managed (23-38 individuals per family) reduced the statistical power for detecting associations at family level. Finally, five families such as F12, F14 and F56 did not show any signal of association and environmental factors might be driving gonadal fate in their offspring.

**Figure 2:**
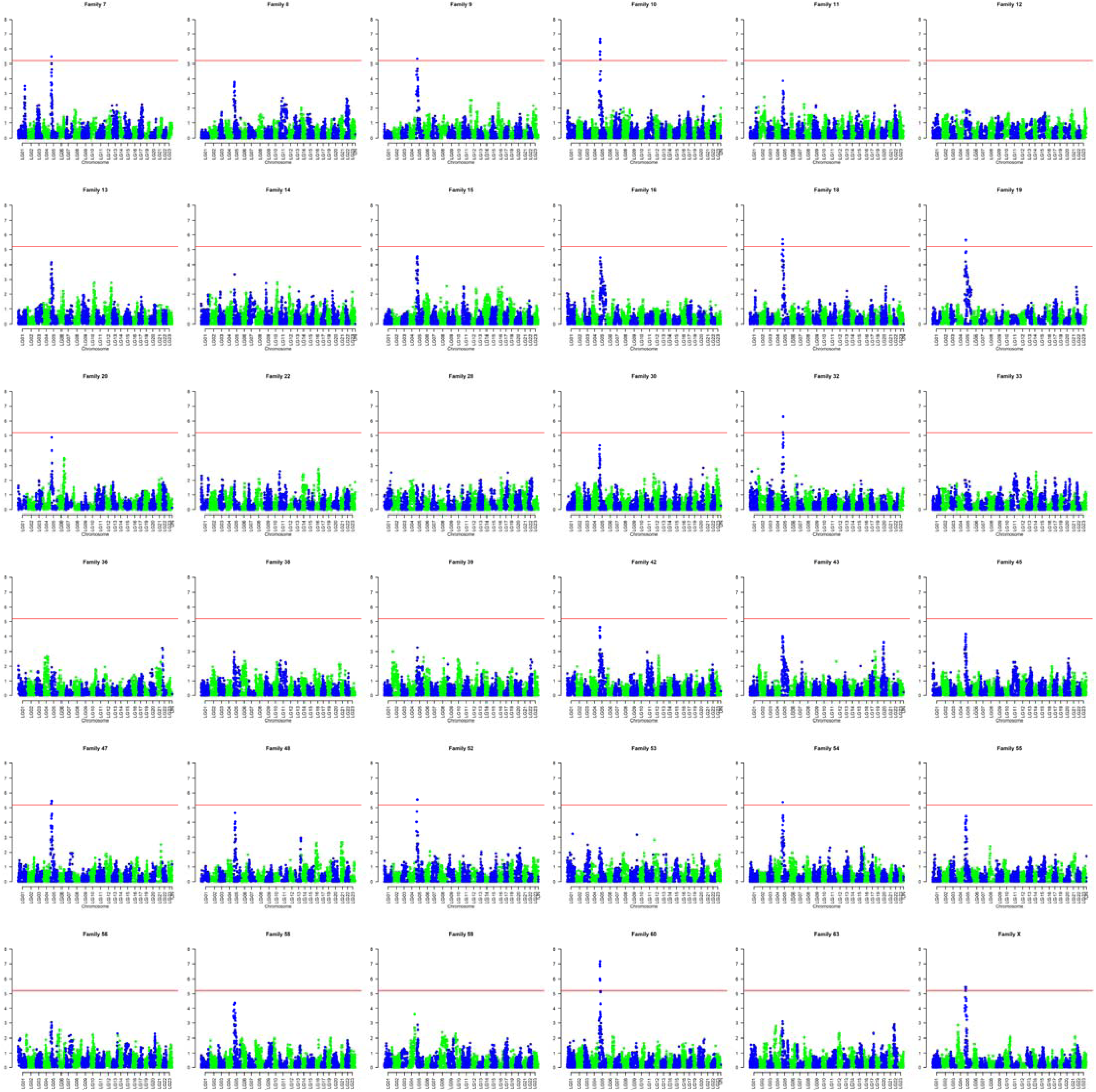
Genome-wide association study (GWAS) on turbot (*Scophthalmus maximus*) sex for each full-sib family in the dataset. The red horizontal bar represents genome-wide significance after Bonferroni correction.

#### Segregation Analysis

A total of 36 full-sib families totalling 1135 sexed offspring and 46 parents were used for a segregation analysis at the major SD region to: i) refine the association of sex with the this region in each family; ii) check for a putative recombination blockage between W and Z chromosomes; and iii) narrow down the region where the SDg is located. Only five out of 36 families (13.9 %) did not show association between phenotypic sex and markers at the major SD region (P > 0.05), while four showed full association (P = 0; 11.1 %), 20 were associated after Bonferroni correction (P < 0.001; 55.6 %) and seven at P < 0.05 (19.4%) (Tables 1 and S3). At individual level, 126 fishes showed a discordant phenotype according to the major SD region marker information in the families significantly associated with sex (P < 0.05; 11.1%), although other 155 fishes pertained to families where no association was detected, thus totalling 281 non-associated individuals (24.8%). The information obtained is compatible with the ZW/ZZ system reported for LG5 in turbot (Martínez et al., 2009), being the mother the responsible for the sex of their offspring, and fully concordant with the GWAS evaluation shown in the previous section at family level. A highly significant trend (P < 10^-5^) of genetic females being phenotypic males was detected in those families where sex was associated with LG5 after Bonferroni correction (55 ♀ → ♂ *vs* 13 ♂ → ♀; Table 1). Moreover, the lowest variance in sex ratio among families was detected in those groups where the SD region showed the highest influence on phenotypic sex (Table 2). Finally, we took advantage of HS families to check if the weight of the SD region on phenotypic sex was consistent across families sharing the same mother (15 HS families) (Table 1). No correlation was detected in the percentage of genetic/phenotypic discordances in the family pairs tested, thus suggesting that the weight of LG5 on SD seemed to depend more on the specific genetic and environmental background of each family than on the genotype of the mother at LG5.

**Table 2:** Average proportion of females in its variance in the groups of families showing different association degree with the main SD region in turbot. N: no association; S: significant association (p < 0,05); B: association after Bonferroni correction (P < 0,001); F: full association (no discordances).

|  | average | variance |
| --- | --- | --- |
| **N** | 58,5 | 385.9 |
| **S** | 45,3 | 619.5 |
| **B** | 40,9 | 110.3 |
| **F** | 45,6 | 57,8 |

A crossing-over map was represented at the proximal region of LG5 where the SD region is located (6 Mb; Figueras et al., 2016) after evaluating the 251 recombinants detected between the W and Z chromosomes of the mother in all families (Table 1; Table S3). Among these, we selected 121 consistent recombinants (at least two markers involved) where the crossing-over point could be delimited within a region < 300 kb. Crossing-over was strongly impeded at the proximal region of LG5 supporting the positional effect of centromeric heterochromatin on the adjacent genomic region (Figure 3). Moreover, crossovers were distributed throughout the rest of the region evaluated, but, interestingly, crossing-over was blocked (or nearly blocked) around the most interesting region, where the genes *sox2*, *dnajc19* and *fxr1* are located. This was not a matter of lower informativeness of markers in that region, since nine families with pairs of informative markers covering this region did not show any signal of recombination between Z and W chromosomes.

**Figure 3:**
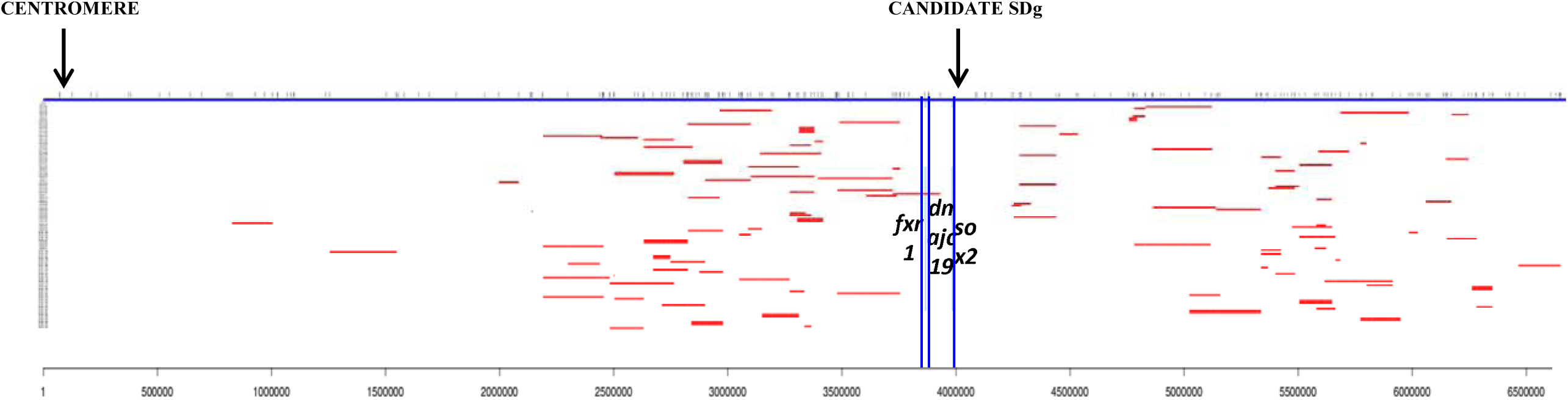
Crossing-over map between Z and W chromosomes (red bars) delimited using the closest informative markers at a distance < 300 kb through recombinant offspring evaluation in 36 families of *S*. *maximus*. The blue upper bar shows the scaled position of SNPs included in the eight scaffolds located at the proximal region of LG5 (∼ 6 Mb). The blue light vertical bars indicate the position of the three candidate genes at the SD region (*fxr1*, *dnajc19* and *sox2*). The scale to the left indicates the code recombinants detected.

Finally, the segregation analysis enabled to narrow down the region where the SDg is located in turbot using the most confident set of informative families (chosen among those showing association after Bonferroni correction) (Figure 4; Table S3). Twelve families were selected for this analysis (F7, F9, F10, F18, F19, F32, F42, F47, F52, F54, F60 and FX) and the smallest region flanked by recombinants at each family was established following conservative criteria, to say, at least two recombinants (see for instance offspring ID 12 and 462 of family 10; Table S3) or the second best recombinant (see offspring ID 240 and 584 of the same family) were used to define the left or right edges of the region containing the SDg in each family. The minimum overlapping region combining the information of all families where the SDg should be located was between 3,733,262 bp and 4,129,527 bp (∼ 365,330 bp; Figure 4). However, considering the presence of adjacent non-informative SNPs at the edges of each recombinant, we decided to be more conservative and took the next informative marker to establish the limit, so expanding the overlapping region across families to 961,390 kb (Figure 4).

**Figure 4:**
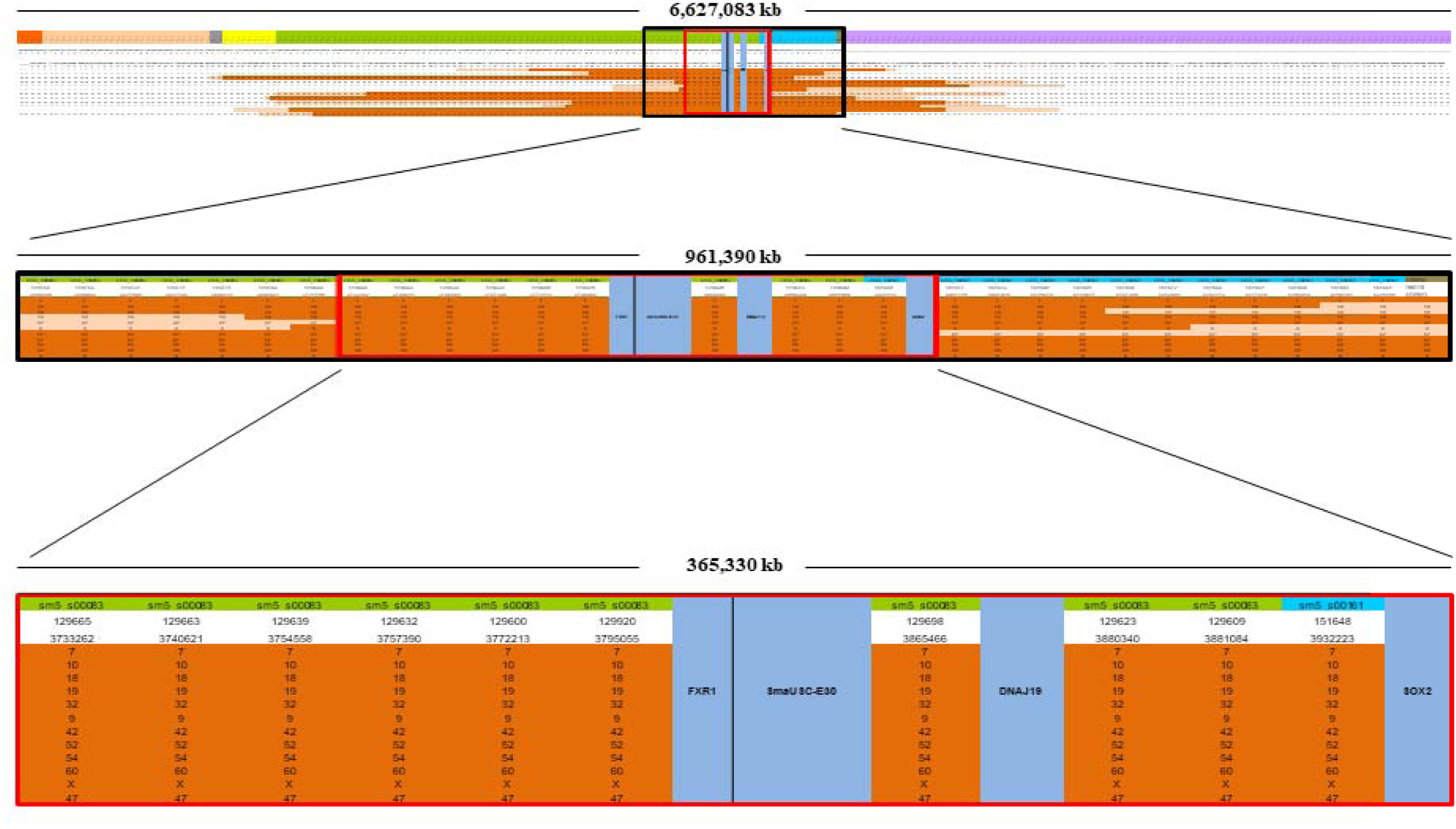
Representation of the segregation analysis at LG5 in the 12 most informative and consistent turbot families to narrow down the genomic region where the SDg is located. Colors in the upper row of each chart show the different scaffolds of turbot genome at LG5 (Maroso et al., 2018). A progressive zoom is illustrated from the whole region analyzed atLG5 (> 6 Mb; upper chart) through the most conservative estimation of SDg location down to the narrowest less conservative estimation (bottom). Brown color indicate the least conservative estimation of the SDg location at each family, while the cream color indicates the region where the most informative crossover/s should have occurred. The black and red squares at the top indicate the minimum overlapping region for each estimation. The three genes and the SmaUSC-E30 marker located in that region of the turbot genome are shown.

### Reassembling the major SD region through hybrid long-/short-read resequencing: diagnostic differences between the W and Z chromsomes

To merge the two main scaffolds (83 and 161; Figure 1) constituting the turbot SD region and to have a confident reference to look for diagnostic differences between the W and Z chromosomes, a superfemale (WW) and a male (ZZ) were sequenced and assembled combining long-read Nanopore and short-read Illumina technologies.

Long-read Nanopore sequencing of the superfemale rendered 848,617 reads with an average length of 6,269 bp (longest read: 84,308 bp) totalling 5,3 Gbp and representing 9.4x coverage of the turbot genome (565 Mb; Figueras et al., 2016); the male sequencing produced 630,136 reads with an average length of 9,197 bp (longest read: 177,236 bp) totalling 5,8 Gbp representing 10.2x coverage. The error rate of Nanopore sequencing was evaluated by matching all reads against the coding regions of the turbot genome and 90.4 % homology was observed on average, thus within the lowest error rate range for this technology (Lima et al., 2019).

Nanopore and Illumina sequencing were used to refine the assembly of the turbot SD region delimited through segregation analysis (961.4 kb; Figure 3b). All Nanopore and Illumina sequences matching to that 961.4 kb region were retrieved for assembly using the last version of the turbot genome (Maroso et al., 2018). A hybrid assembly was obtained separately for the W and Z chromosomes using Nanopore assembly as the backbone, which was then curated using the Illumina 150 bp PE reads of five WW females and five males, respectively. Both the superfemale and male hybrid assemblies covered > 95% of the delimited SD region including the gap between the scaffolds 83 and 161, the region where the strongest sex association was detected (Figure S2). Further, the combination of the W and Z assemblies covered the whole 961.4 Mb region of the turbot genome (Figueras et al., 2016). It should be noted that the turbot genome was constructed using a female (Figueras et al., 2016), and thus, misassembling at the differential region between the W and Z chromosomes could occur. No evidences for duplication of the candidate genes reported in turbot by Taboada et al (2014) or structural reorganizations, such as inversions, were detected.

The five WW females and males re-sequenced using Illumina 150 bp PE reads were separately aligned against the W and Z hybrid assemblies and alignments were inspected for diagnostic differences between WW females and males using the graphic viewer IGV 2.4.10 (Robinson et al., 2011). Only two SNPs showed a diagnostic association with W and Z chromosomes across the inspected SD region (SNP.1 and SNP.3). These SNPs, fixed for alternative allelic variants in Z and W chromosomes, were located between the genes *dnajc19* and *fxr1* (Figure 5), surrounding the SmaUSC-E30 microsatellite used for sexing turbot (Martínez et al., 2014). A SNaPshot assay was designed to evaluate the consistency of this association at species level in a sample of 92 individuals, 46 males and 46 WW females, belonging to 11 unrelated families of a turbot breeding program were genotyped for these two SNPs (Table S4). Genetic differentiation between males and WW females for these two SNPs was very high and significant (F_ST_ = 0.576 and 0.500 for SNP.1 and SNP.3, respectively; P = 0) and extreme linkage disequilibrium was detected between them (P = 0). Most males showed an AAGG genotype while most WW females were GGAA suggesting an AG haplotype for the Z chromosome and GA haplotype for the W chromosome (Table S5); seven individuals showed an AGAG genotype congruent with a ZW constitution and five WW females showed an unexpected GGAG genotype. These WW females pertained to a single family suggesting that other low frequency haplotype might exist at the W chromosome. All in all, SNP.3 was compatible with a diagnostic marker between Z and W chromosomes assuming the ZW constitution of the seven aforementioned individuals.

**Figure 5:**
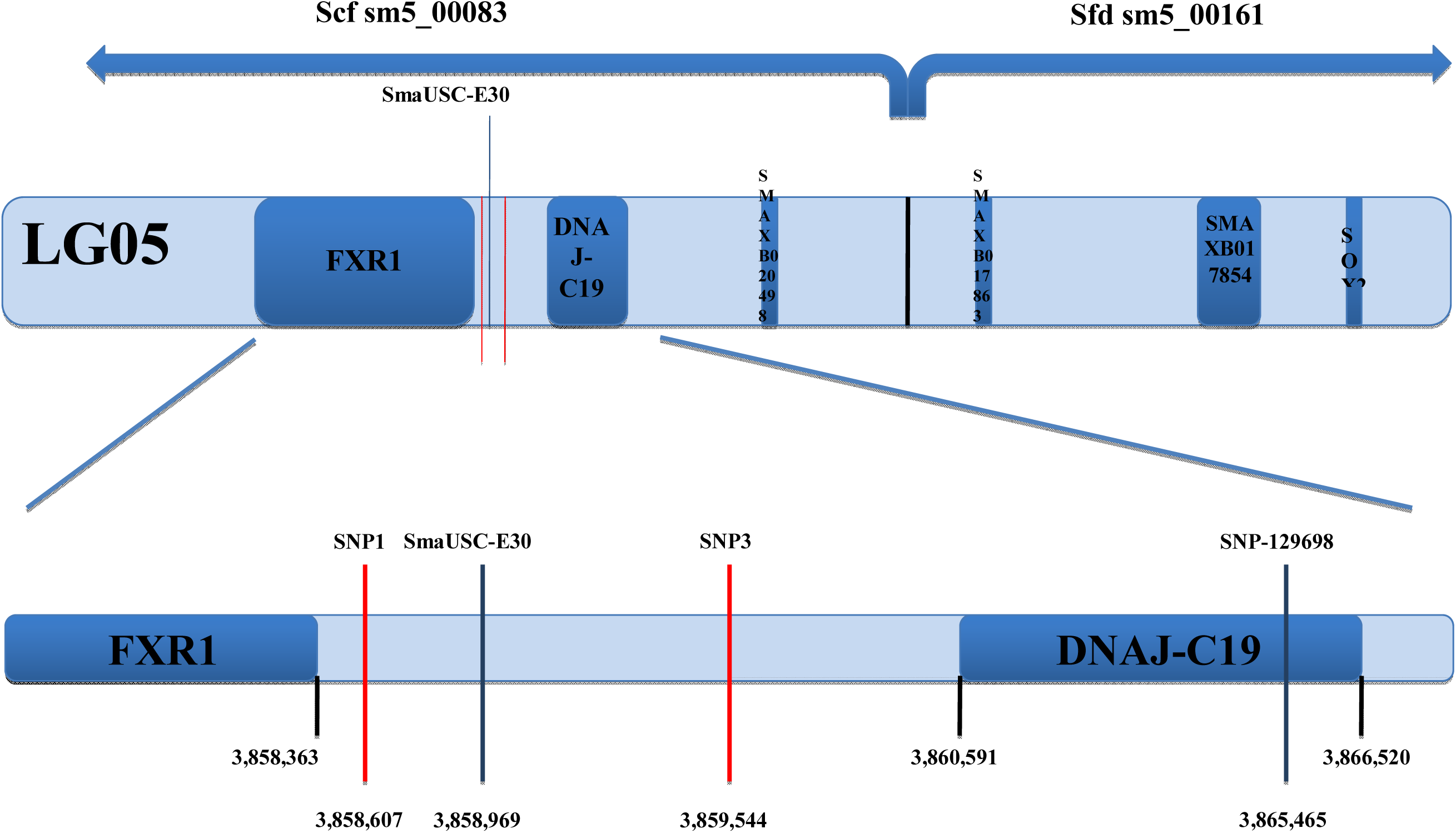
Scheme of the turbot SD region including the two SNPs discriminating the W and Z chromosomes surrounding the SmaUSC-E30 sex-associated microsatellite (Martínez et al., 2009) and the annotated genes and ORFs. The two scaffolds of the turbot genome at that region are shown in the upper part; below a zoom including the positions of the main reference points.

### Functional analysis: comparative gene expression between sexes along gonad differentiation (GD)

#### Identification of active genes along gonad development and differentiation

To identify the genes expressed along the gonad differentiation period at the major SD region in turbot, ribodepleted RNA libraries obtained from a pool of larvae, a pool of male gonads and a pool of female gonads at developmental stages from 5 dpf until 90 dpf were sequenced. A total of 24 genes were expressed along that period in the delimited 961.4 kb region (Figure 6; Table S6). Among the three previously reported candidate genes in the region, *dnajc19, fxr1* and *sox2* (Taboada et al., 2014), only the two latter were expressed. Kininogen-1 (*kng-1*), complement factor H (*cfh*) and Fragile X mental retardation syndrome-related protein 1 (*fxr1*) were the highest expressed genes, while the lowest one was *sox2ot* (see below). None of the genes in the region showed obvious signs of differential expression between the male and female pools obtained from samples encompassing the whole critical period of gonad differentiation (between 35 and 90 dph; Figure 7), an expected result if the critical development window where differential expression between sexes is short. Moreover, a few genes showed important differences between larvae and more advanced developmental stages, including *kng-1, cfh* or *sox2*.

**Figure 6:**
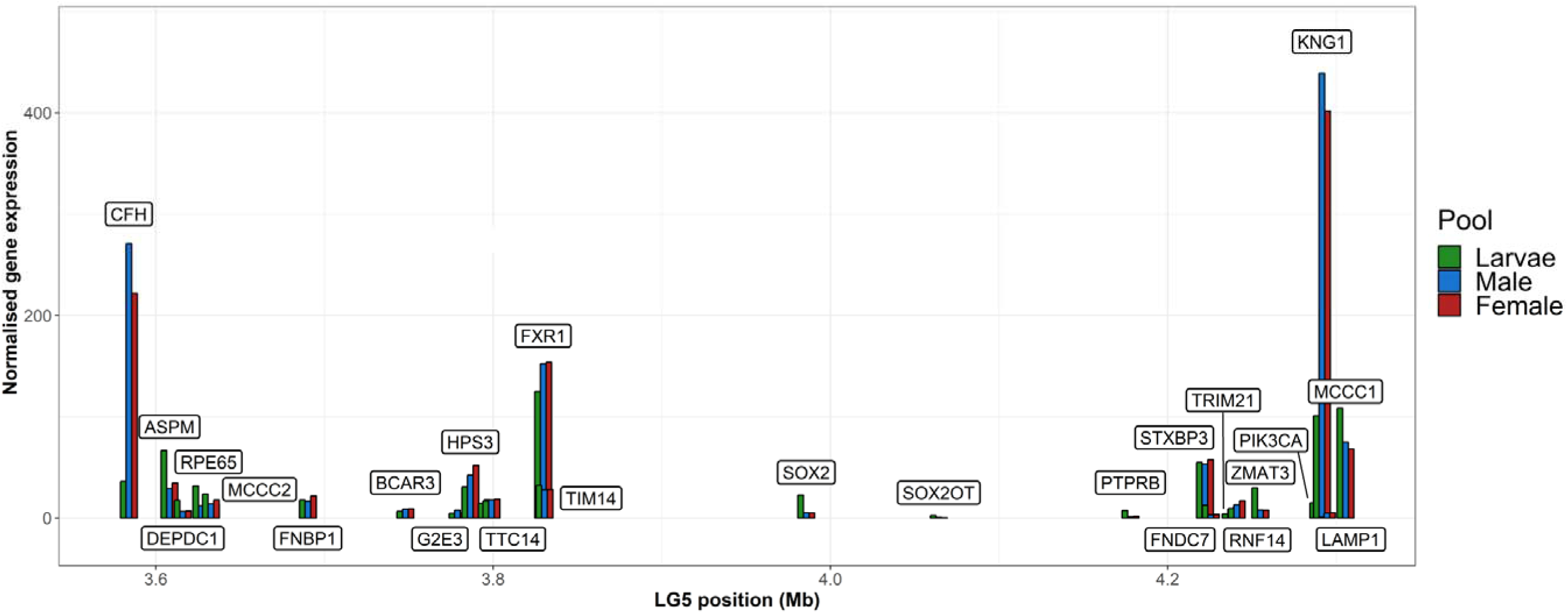
Normalized expression of the transcripts detected in the SD region delimited through segregation analysis (∼ 1 Mb) across the gonad differentiation period (< 90 dph) in pools of larvae, females and males sexed using the SmaUSC-E30 marker (Martínez et al., 2014).

**Figure 7:**
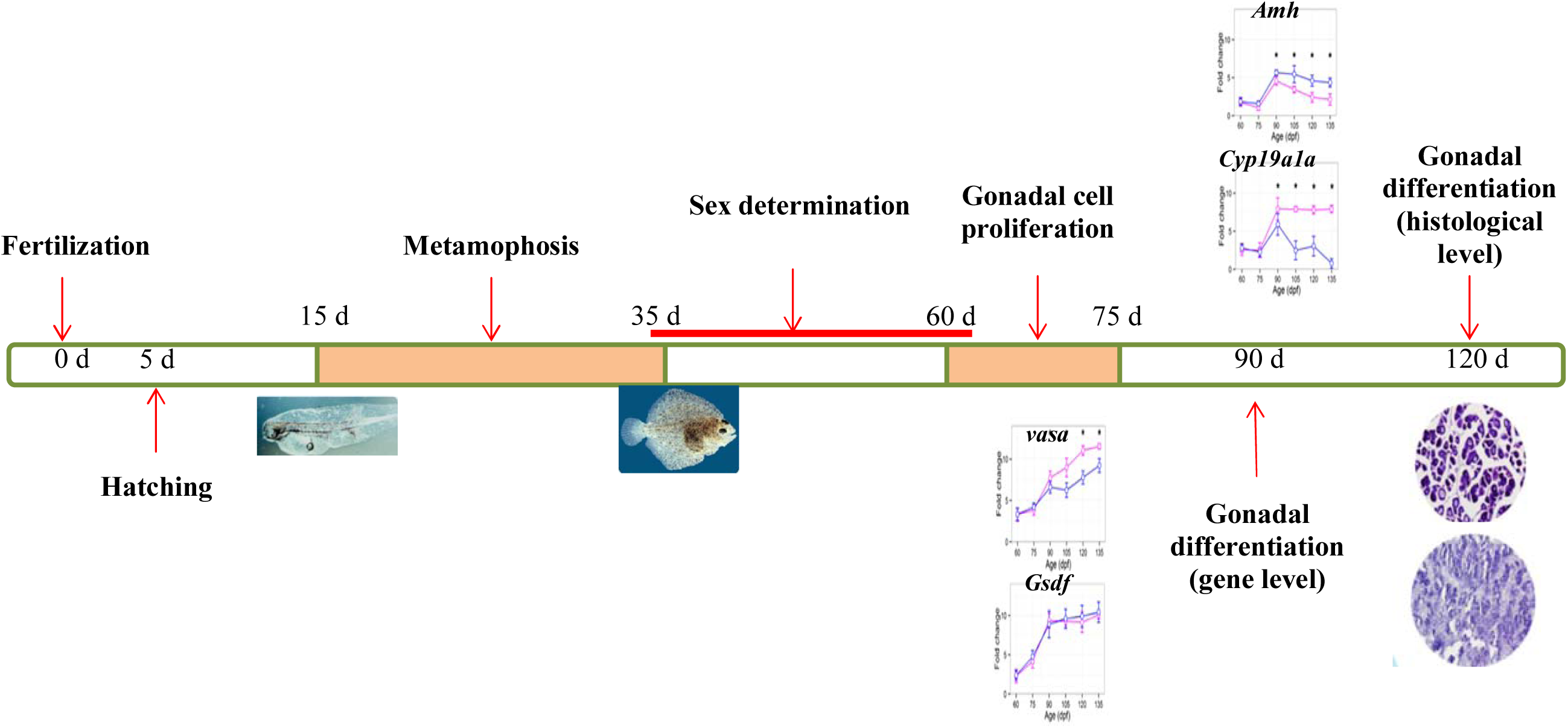
Main development and genetic events across the critical period of gonad differentiation in turbot (qPCR graphs: males in blue and females in pink from Robledo et al., 2015).

A long-non coding RNA encompassing 78,309 bp was found in the region were *sox2* (2,557 bp) is located and it was annotated as *sox2* overlapping transcript (*sox2ot*). This gene, conserved across vertebrates (Amaral et al., 2009), includes the *sox2* gene in one of its introns. Within this lncRNA, a splicing variant corresponding to a short non-coding RNA transcript showed a faint expression along the gonad differentiation period (ncRNA) both in males and females. Interestingly, a ∼ 250 bp specific region of this ncRNA was complementary to a region of the *sox2* mRNA 3’ UTR (Figure S3).

#### qPCR of candidate genes along the critical period of gonad differentiation

Gene expression of candidate genes was individually evaluated by qPCR in males and females along gonad development, particularly between 30 and 90 dph, to say, from weaning until the time where gonad differentiation was established at genetic level (up-regulation of *amh* and *cyp19a1a* in males and females, respectively; Figure 7; Robledo et al., 2015). This is the period where gonads can be dissected with confidence for gene expression evaluation, but specially, it is the development period where SD likely occurs in turbot, after metamorphosis but before gonads are differentiated at functional level. No expression differences were detected for *fxr1*, *dnajc19* and the *sox2ot* ncRNA between males and females, but a consistent pattern of increasing expression was observed for *sox2* in females between 50 and 55 dpf (Figure 8; Table S7). Indeed, significant differences were detected at 55 dpf by considering the information of both amplicons (t-test related samples = 5.153, P < 0.004) and it was even higher when the 50-55 dpf window information was merged (t-test related samples = 6.848, P < 0.0005).

**Figure 8:**
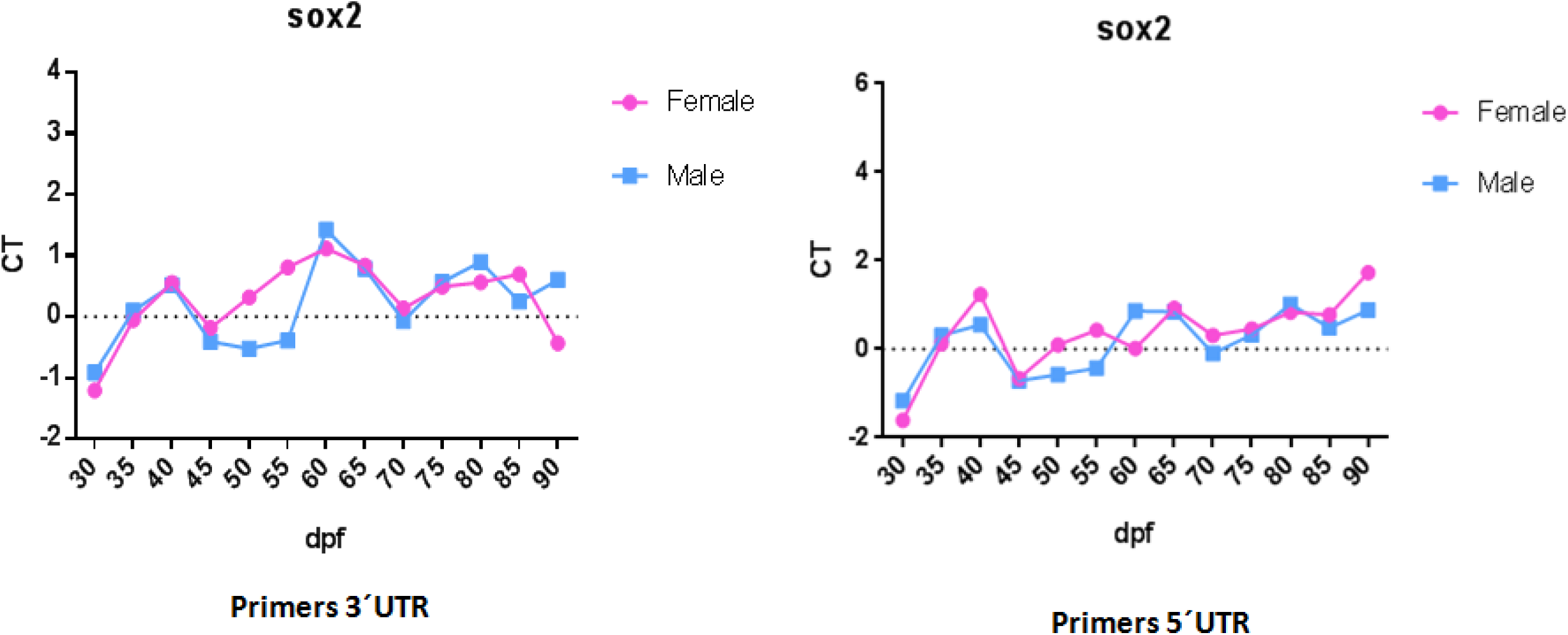
qPCR analysis of *sox2* gene (primers 3′ and 5′UTR) in gonad of male and female between 30 to 90 dpf using ΔCT method with the ribosomal protein S4 (RPS4) and ubiquitin (UBQ) as reference genes. An increasing expression is observed between 50 and 55 dpf in females.

## DISCUSSION

Elucidating how sex is developed and controlled in vertebrates in general and in fish in particular is a flourishing area of research from evolutionary and development biology perspectives. The high turnover of SD systems and the small differentiation of sex chromosomes in fish, even in taxa with conserved SD systems, suggest a different scenario to that previously reported in mammals and birds. Additionally, sexual dimorphism affects important aquaculture traits such as growth rate, flesh quality or caviar production, among others, and thus, controlling sex ratio is a key issue for producers (Martínez et al., 2014). For this study, we went beyond the important information previously gathered on sex determination (SD) and gonad differentiation (GD) in turbot by combining genomics and classical genetics approaches, as well as using functional and statistical association methodologies. A large number of families and individuals, as well as a careful selection of biological material was used to ensure the genetic constitution of fishes considering the complex architecture of SD reported in turbot (Taboada et al., 2019). We got a comprehensive picture of the complex mechanism underlying SD in this species and framed this information within teleost SD evolution. In particular, our work represents a contribution to the knowledge of the origin and evolution of “young sex chromosome pairs” (Charlesworth, 2019), still very unknown unlike the most studied mammalian, avian and *Drosophila* systems, which represent “old sex chromosomes” with specific genetic features: size heteromorphism, specialized gene content, reduced recombination and degeneration (Abbott et al, 2017).

### The main sex determining gene (SDg) of turbot

The extensive genome-wide screening performed with 18,214 SNPs (1 SNP / 28 kb; 565 Mb turbot genome; Figueras et al., 2016) in combination with a classical segregation analysis carried out in 36 families (44 parents and 1,135 sexed offspring) confirmed that SD of turbot mainly relies on a small region at the proximal end of LG5 and follows a ZW/ZZ pattern of inheritance (Haffray et al., 2009; Martínez et al., 2009; Taboada et al., 2019). However, unlike previous reports, here we gathered enough evidences to suggest *sox2* as the major SDg in this species: i) it is located in the narrow window established after segregation analysis and surrounded by the two most significant SNPs identified by GWAS; and ii) it is the only gene among the transcripts expressed along GD in the delimited SD region that showed a differential expression between males and females within a short five days window (50-55 dpf), just before the first signals of germ cell proliferation (up-regulation of *gsdf1* and *vasa* at 60-65 dpf; Robledo et al., 2015). This gene had been previously discarded as the SDg of turbot because no diagnostic SNPs could be identified in the coding region in a large sample of males and females (Taboada et al., 2014). Despite this still holds, data suggest that the responsible mutation for differential expression of *sox2* between sexes could be located on a regulatory element. In this regard, it should be noted that the lncRNA *sox2ot*, a regulatory gene of *sox2* (Amaral et al, 2009; Shahryari el al., 2015; Messemaker et al., 2018), encompasses the most associated SD region including *sox2* in one of its introns. Further, a splicing variant of *sox2ot* showed full complementarity to *sox2* turbot 3’ UTR, thus suggesting a putative post-transcriptional mechanism controlling *sox2* activity. Although lncRNAs are poorly annotated, they have been involved in the regulation of genes related to reproduction and particularly to SD in different species. In *Drosophila* the master *Sxl* gene is regulated by a lncRNA acting in a complex interplay network (Mulvey et al., 2014) and in the crustacean *Dhapnia magna* a lncRNA regulates the male specific expression of the *dsx1* DM domain producing males in response to environmental stimuli (Kato et al., 2018). Moreover, the only diagnostic variant detected between Z and W chromosomes in our study (SNP.3), despite being far from *sox2*, might be part of a regulatory element, since *sox*2 enhancers have been identified within a 200 kb region surrounding this gene (Zhou et al., 2014). Recently, *sox2* has been identified as the candidate SDg in two other aquaculture species, the Zhikong scallop *Chlamys varia* (Liang et al., 2019) and the black tiger shrimp *Penaeus monodon* (Guo et al., 2019).

### Turbot SD as a complex trait

Our data strongly suggest a complex genetic architecture of turbot SD, supporting previous information (Martínez et al., 2014; Taboada et al., 2019). Despite the major SD region drives GD in nearly 80% of the fish, and close to 90% of the families show a significant association between sex and markers in that region, other genetic and environmental factors interact with the main locus to fate the sex of each fish. The amount of data here gathered (1,181 individuals; 36 FS families; 27 HS families) enabled deepening into the different factors involved. Other sex-related QTL had been previously reported in turbot, always as secondary players of the major SD locus in the families analyzed (Martínez et al., 2009; Hermida et al., 2013). We confirmed these observations, but additionally demonstrated that in some families another genomic region different than LG5 would be the main driver for GD. None of the previously reported secondary QTL were identified in the broad family sample analyzed in this study, and those here detected seemed to influence sex in only one or two families. The most consistent one was identified at LG22 which showed signals of association in two families and at chromosome-wide level in the whole dataset. These observations suggest either the presence of rare allelic variants affecting sex in several secondary QTL or, more likely, complex genetic and/or environmental interactions providing scenarios for other genomic regions to drive SD. Our observations would also fit with the stochastic gene expression and development noise on SD suggested by Perrin (2016).

Moreover, environmental factors influencing SD have been described in fish, where high rearing temperatures tend to induce masculinization (Ospina-Alvarez and Piferrer, 2008) but the opposite is still controversial. In the case of turbot, the influence of temperature does not seem to work in a simple way (Haffray et al., 2009), although Robledo et al. (2015) showed a significant trend for female biased ratios at low temperatures. We identified four families where no signals of genomic association were observed, suggesting environmental factors driving SD. Furthermore, the proper weight of the major SD region in each family seemed to not follow a simple additive model, since mothers of HS families showed a variable influence of this region in the families they damed. The existence of different genetic factors and even a polygenic architecture underlying SD in fish have been documented in several species such as zebrafish, European sea bass and tilapia (Vandeputte et al., 2007; Nagabhusana and Misrha, 2016), among others. Even in those species where a master SDg has been reported, deeper studies identified the presence of secondary genetic and environmental factors (Martínez et al., 2014). Data from turbot supports the influence of environmental factors, mainly temperature, but following complex interactions with genetic factors, thus approaching to a complex trait.

### SD in teleosts: some insights from turbot

Knowing the full sequence of sex chromosomes is essential to identify the causative mutations underlying SD and to quantify their differentiation degree in order to understand their evolutionary pattern. Despite the increasing number of genomes sequenced and assembled, the sequences of sex chromosomes Y or W are mostly unknown because the presence of repetitive elements represents a handicap for their assembly and usually require at least twice the depth of a homogametic genome (Tomaszhiewicz et al., 2017). In the case of fish, the early evolutionary stage of sex differentiation and the viability of WW or YY individuals (Devlin and Nagahama, 2002) facilitate this task, especially if the long-read sequencing technologies are combined with the more confident short-read ones to achieve better scaffolding (Tan et al., 2018; Xing et al., 2019). We sequenced and assembled the Z and W chromosomes of turbot using five females (WW) and five males (ZZ) by combining long-read Nanopore and 150 bp PE Illumina methods with enough coverage to detect structural differences, essential to understand the evolution of SD. Further, this strategy also allowed us to detect minor differences that could be related to the causative mutation determining sex in turbot. As previously suggested by Taboada et al. (2014), we confirm that the major SD region of turbot is a young one (Charlesworth, 2019), since no consistent genetic differences between Z and W chromosomes more than in a single nucleotide (SNP.3) could be detected. In this regard, turbot’s would be among the youngest sex chromosome pairs found in fish at a stage similar to that of the pufferfish (*Fugu rubripes*; Kamiya et al., 2012), where the only difference between X and Y chromosomes was a single differential SNP at the promoter of the *amh* receptor. Very recently, a similar picture has been reported in *E*. *lucius*, where a small male-specific insertion containing the SD gene, has been documented (Pan et al., 2019). Although examples of mature SD systems such as those reported in the genus *Characidium* and *Eigenmannia* have been reported (Cioffi et al., 2017), the differential region in most species where the master SDg has been identified comprehends less than 1 Mb (Martínez et al., 2014). It is generally accepted that the evolution from a homomorphic to a heteromorphic sex chromosome pair begins with the suppression of recombination to maintain particular haplotype combinations to avoid sexual conflict (Bull 1983; Charlesworth, 1994, 1996). Our data suggest that crossing-over is precluded in a region of ∼75 kb around *sox2*, mostly embracing the lncRNA *sox2ot*, thus representing the first stratum in the evolution of the sex-chromosome pair of turbot. Taboada et al. (2014) also suggested a transition from an ancestral XY to a new ZW system and indicated a non-complete dominance of the ZW reflected by a significant amount of ZW males in the families analyzed, a fact here confirmed with a much larger amount of data. Finally, we obtained suggestive information about the increasing genetic canalization of SD, characterized by a much higher sex ratio balance, in those families where the LG5 SD region is dominant with regard to those families with secondary genetic or environmental factors involved, which showed more unbalanced sex ratios either towards males or females.

It is worth a reflection on why SD of fish is so unstable in evolutionary terms, where the classical model of two well differentiated sex chromosomes is scarce. The high turnover of SD in fish suggests that different genomic regions can be recruited to replace the previous SD system following the so called “hot-potato” model (Gammendinger and Kocher, 2018), although data strongly suggest the suitability of some genes like *dmY* or *amh*, recruited independently in several fish species to drive sex differentiation (Matsuda et al., 2002; Hattori et al., 2012; Pan et al., 2019). In this regard, *sox2* and the putative involvement of *sox2ot* lncRNA on its regulation, would add a new master gene and a novel regulatory mechanism confirming the multiple options to drive sex across the fish genomes. As shown in our gene expression analysis, *sox2* is very active across larval development, likely governing important functions related to neurosensory development as reported in other species including fish (Gou et al., 2018; Steevens et al., 2019). Since no duplication of *sox2* was detected in our hybrid SD assembly, data suggest that the complex regulation of this gene, involving a lncRNA and several long-distant regulatory elements, would have been harnessed for gonad differentiation at a later developmental window through its up-regulation in females. It should be noted that the involvement of *sox2* in different traits including GD could underlie a pleiotropic explanation on the origin of the SD of turbot, as has been hypothesized (Gammendinger and Kocher, 2018).

### Applications for turbot farming

One of the goals of this work was to find a reliable genetic marker to be applied for sexing by turbot industry. It should be noted that in turbot no morphological dimorphism exists between sexes prior to sexual maturity, and therefore sex cannot be identified at juvenile stages (Martínez et al., 2016). The marker USC-E30, a microsatellite used as sexing tool under a Spanish patent (Ref. number: 2354343), although useful and being applied by industry to obtain all female populations, has the drawback of the high and recurrent mutation rate of microsatellites, which obligates to establish associations within each family and to genotype relatives of the parents to identify the specific association. In this sense, the SNP markers associated to Z and W chromosomes in our study will provide a more confident and straightforward tool for precocious sexing aimed to improve management of breeding programs and to obtain all-female populations by industry.

### Conclusions

The turbot SD system is among the youngest reported in fish and only a single differential SNP could be confidently identified between Z and W chromosomes. Recombination between Z and W chromosomes seems to be suppressed at a short stretch of ∼ 75 kb that could represent the first differentiation stratum. *Sox2*, a new gene of the transcription factor *sox* family to be added to the large list of fish SDg, appears to be the master SDg of turbot and it is located within an intron of the lncRNA *sox2ot*. This lncRNA may be involved in the regulation of *sox2* since the homology detected between a *sox2ot* splicing variant and its 3’ UTR. Other genomic regions were associated with sex, particularly at LG22, being the most associated region in a subset of families. However, these secondary factors seem to be of small effect and dispersed at different genomic regions. Furthermore, environmental factors could be the main driver fating sex in a number of families. The major SD region at LG5 appears to better canalize sex ratio in families providing an improved mechanism to avoid demographic uncertainty. The weight of this region on SD does not seem to follow an additive model, and again, interactions are needed to explain its role. The information here gathered will be important for obtaining all-female populations by industry using the differential SNP identified between Z and W chromosomes at species level.

## MATERIALS AND METHODS

### Sampling

Different sets of samples were used for the different methodological approaches carried out to identify the sex determining gene (SDg) of turbot and to provide functional evidences of its role along the critical stages of gonadal differentiation: i) 1181 fish (between 133 and 217 days after hatching – dph) pertaining to 36 full-sib families (1135 offspring and 46 parents) were sexed by histology and genotyped using 2bRAD-Seq (Wang et al., 2012) for 18,214 SNPs to look for association between sex and genetic markers at population and family level through genome wide association studies (GWAS). This biological material was also used for a segregation analysis that enabled to narrow down the region were the SDg is located and to establish a recombination map at the SD region. These 1181 fishes came from an experiment for resistance to scuticociliatosis performed at CETGA (Centro Tecnológico Gallego de Acuicultura) facilities within the FISHBOOST project (KBBE-2013-7-613611); ii) Five males (ZZ) and five WW females (WW) were used to refine the assembling of the turbot SD region by combining 150 bp paired-end (PE) Next-seq Illumina sequencing and the long-read Nanopore technology to identify genetic differences between the W and the Z chromosomes, and eventually, the causative variant responsible for SD. WW females were obtained following a three-generation pedigree as described by Martínez et al. (2014); briefly, this protocol combines the use of methyltestosterone to obtain neo-males and the use of the marker SmaUSC-E30 to predict the genetic sex (Martínez et al., 2009) in neomales ZW and WW females WW. The five ZZ males and the five WW females pertained to different unrelated families and were obtained from the breeding program of a turbot company; iii) Larvae or fry from 5 to 90 dph (18 sampling points) were pooled for RNA-seq analysis (six individuals per sampling point) to identify transcripts within the SD region expressed from hatching until the gonadal fate is established (90 dph; Robledo et al., 2015). These samples were obtained at the IEO (Instituto Español de Oceanografía) of Vigo using 10 families of unrelated parents; thirteen of those samples, including three males and three females per sampling point, were further used for individual qPCR gene expression analysis of candidate SD genes along the critical period of turbot gonad differentiation.

### High-throughput SNP genotyping

A 2b-RAD method (Wang et al., 2012) was applied to genotype 18,214 SNPs in the 1181 fish used for GWAS and segregation analysis as reported by Maroso et al. (2018). Briefly, SNPs were filtered out according to SNP call-rate < 0.8, individual call-rate < 0.8, identity-by-state > 0.95 (both individuals removed) and minor allele frequency (MAF) < 0.01. The filtering done enabled to obtain a very consistent set of SNPs showing a genotyping error > 0.4 % estimated using family data (Maroso et al., 2018).

### Pinpointing the turbot SDg: genome-wide association study (GWAS) and segregation analysis

GWAS was performed using GenABEL R package (Aulchenko et al., 2007) by applying the mmscore function (Chen and Abecasis, 2007), which accounts for the relatedness between individuals using a genomic kinship matrix. Significance thresholds were calculated using a genome-wide Bonferroni correction where significance was defined as 0.05 divided by the number of informative SNPs (Duggal et al., 2008). GWAS were performed using the whole population data as well as within each family.

A segregation analysis, which complemented GWAS, was done to identify the minimum genomic region where the SDg is located and to gather additional information on the genetic components and properties of the SD region. For this, only those informative families showing a highly significant sex association with markers at the SD region were used. The availability of parents and offspring genotypes and the high SNP density used (1 SNP/ 28 kb) enabled to infer the haplotypes constituting the Z and W chromosomes of the mother within each family at LG5, the linkage group where the major SD region is located (Martínez et al., 2009). Then, those offspring showing crossovers between the Z and W chromosomes were analyzed and its sex recorded to check if the fraction of the W chromosome inherited from the mother did not include (males) or include (females) the SDg. Finally, information of the families was combined to delimit the region where the SDg would be located. Additionally, this information was used to identify those individuals that, despite having inherited the whole W or Z chromosomes from the mother, showed an unexpected sex phenotype, to say, a ZW male or a ZZ female, thus suggesting other genetic or environmental factors involved. Finally, a recombination map of the SD region was constructed using the aforementioned information to identify the distribution of cross-overs and to check for a putative recombination blockage between the W and Z chromosomes.

### Improving the assembly of the SD region: searching for the SD causal variant

Five males and five WW females were used to refine the assembly of the turbot SD region using both 150 bp Illumina PE sequencing reads and long-read Nanopore sequencing. Then, we looked for consistent genetic differences between the W and the Z chromosomes, including putative reorganizations or duplications, which could be related to SD in turbot. These 10 individuals were carefully selected using families where information on relatives was available and that had been genotyped for several markers at the SD region to ensure the ZZ and WW constitution of males and WW females, respectively.

#### Nanopore sequencing

DNA from one male and one superfemale was extracted using the MagAttract HMW DNA Kit, which ensures reproducible isolation of genomic DNA >150 kb. The procedure comprised four simple steps: lysate, binding, washing and elution. Following sample lysis, the DNA was bound to the surface of magnetic beads; during the washing steps, contaminants and PCR inhibitors were removed and pure high-molecular-weight DNA was eluted in Buffer AE. Two runs, one for the male and the other for the superfemale, at an estimated 10x coverage, were performed on a Minion Oxford Nanopore sequencer at CNAG (Centre Nacional de Análisis Genómico, Barcelona).

#### Illumina sequencing

Genomic DNA of five males and five WW females was extracted from fin clips using SSTNE buffer (a TNE buffer modified by adding spermidine and spermine) and a standard NaCl isopropanol precipitation (Cruz et al., 2017). Barcoded libraries of 150 bp PE were constructed for each individual and subsequently sequenced in a Next-seq Illumina machine at an estimated coverage of 20x per individual. The quality of the sequencing was assessed using FastQC v.0.11.5 (http://www.bioinformatics.babraham.ac.uk/projects/fastqc/). Quality filtering and removal of residual adaptor sequences was conducted on read pairs using Trimmomatic v.0.36 (Bolger et al., 2004). Specifically, Illumina adaptors were clipped from the reads, leading and trailing bases with a Phred score < 20 were removed, and the read trimmed if the sliding window average Phred score over four bases was < 20. Only reads where both pairs were longer than 36 bp post-filtering were retained. Filtered reads were mapped to the most recent turbot genome assembly (ASM318616v1; Genbank accession GCA_003186165.1; Maroso et al., 2018) using Burrows-Wheeler aligner v.0.7.8 BWA-MEM algorithm (Li, 2013). Pileup files describing the base-pair information at each genomic position were generated from the alignment files using the mpileup function of samtools v1.6 (Li et al., 2009), discarding those aligned reads with a mapping quality < 30 and those bases with a Phred score < 30.

#### Illumina-Nanopore hybrid assembly

A *de novo* hybrid assembly with Illumina and Nanopore reads was carried out at the delimited SD region (see segregation analysis) using those reads matching at LG5 in that region using the last version of the turbot genome assembly (Maroso et al., 2018). Assembly was performed with the selected reads following the instructions provided by Oxford Nanopore (https://nanoporetech.com/resource-centre/hybrid-assembly-pipeline-released-using-canu-racon-and-pilon); briefly, assembly of Nanopore reads was done using Canu assembler (Koren et al., 2017), resulting contigs were polished with the module Racon (Vaser et al., 2017), and finally, the assembly was curated using the Pilon tool (Walker et al., 2014) by adding the Illumina reads.

#### Genetic differences between the W and Z chromosomes: identification and validation of the causal SD polymorphism

The alignment of Illumina reads from five males (ZZ) and five WW females (WW) to the Z and W hybrid chromosome assemblies enabled the identification of sex chromosomes specific genetic variants. The small sample size and the limited coverage of this re-sequencing evaluation suggested a further validation on a larger sample of males and females. For this, a sample of 46 males and 45 WW females coming from 10 unrelated families was used to check for this association at species level using either a PCR on agarose gel for large indels or a SNaPshot protocol for SNPs. Primers for PCR amplification were designed with Primer 3 (http://bioinfo.ut.ee/primer3/) and amplifications were checked on 1 % agarose gels. Additionally, an internal primer, either forward or reverse, was designed for the SNPs mini-sequencing reaction of the SNaPshot protocol. SNP genotyping was finally done in an ABI3730 automatic sequencer (Applied Biosystems). Genetic differentiation between Z and W chromosomes was investigated for specific markers using the relative coefficient of differentiation (F_ST_) and their pairwise linkage disequilibrium using genotype association exact tests both implemented in GENEPOP 4.0 (Rousset, 2008).

### RNA sequencing (RNA-seq)

RNA extraction was performed using the RNeasy mini kit (Qiagen) with DNase treatment and RNA quality and quantity were evaluated in a Bioanalyzer (Bonsai Technologies) and in a NanoDrop® ND-1000 spectrophotometer (NanoDrop® Technologies Inc), respectively. RNA samples from whole larvae or male and female fry gonads across the gonad formation and differentiation period (1 to 90 dph) were evenly pooled for sequencing 100 bp PE reads on an Illumina HiSeq 2000 at the Wellcome Trust Centre for Human Genetics, Oxford. Male and female gonads were obtained from 35 dpf until 90 dpf, when gonads could be identified and extracted, and the genetic sex of each individual established using the SmaUSC-E30 marker (Martínez et al., 2014). The quality of the sequencing was assessed using FastQC v.0.11.5 (http://www.bioinformatics.babraham.ac.uk/projects/fastqc/). Quality filtering and removal of residual adaptor sequences was conducted on read pairs using Trimmomatic v.0.36 (Bolger et al., 2004). Specifically, Illumina adaptors were clipped from the reads, leading and trailing bases with a Phred score < 20 were removed and the read trimmed if the sliding window average Phred score over four bases was < 20. Only reads where both pairs were > 36 bp post-filtering were retained. Filtered reads were mapped to the most recent turbot genome assembly (ASM318616v1; Genbank accession GCA_003186165.1; Maroso et al., 2018) using STAR v.2.5.2b (Dobin et al., 2013), the maximum number of mismatches for each read pair was set to 10 % of trimmed read length, and minimum and maximum intron lengths were set to 20 bases and 1 Mb respectively. STAR alignment files were used to reconstruct the turbot transcriptome using Cufflinks v.2.2.1 (Trapnell et al., 2010). Transcript abundance was quantified and normalized using kallisto v0.44.0 (Bray et al., 2016).

### Real-Time PCR

The RNA samples across the gonad differentiation period (1 to 90 dph) were individually analysed by qPCR to test gene expression on candidate genes. As outlined above, male and female fry were genotyped using the SmaUSC-E30 marker. Primers for candidate genes at the major SD region identified by their genomic position and their expression on the GD period were designed for qPCR using the Primer 3 software. Reactions were performed using a qPCR Master Mix Plus for SYBR Green I No ROX (Eurogenetec) following the manufacturer instructions, and qPCR was carried out on a MX3005P (Agilent Technologies). Analyses were performed using the MxPro software (Agilent). The ΔCT method was used to estimate expression taking the ribosomal protein S4 (RPS4) and ubiquitin (UBQ) as reference genes. These two genes had been previously validated for qPCR in turbot gonads by Robledo et al. (2014). Two technical replicates were included for each sample. T-tests were used to determine significant differences between sexes.

## Acknowledgements

This work was supported by the Spanish Ministry of Economy and Competitiveness, Grant: AGL2014-57065-R. We are grateful for the biological material provided by the FISHBOOST EU project (No. 613611), the Cluster de la Acuicultura de Galicia (CETGA) and the Instituto de Oceanografía (IEO) de Vigo. We acknowledge the support provided by Centro de Supercomputaciòn de Galicia (CESGA) in the sequences analysis. We wish to appreciate the technical assistance by Lucía Insua and Sonia Gómez and the support for SNP genotyping provided by Dr. Manuel Vera and Dr. Francesco Maroso.

## References

Abbott JK, Norden AK, Hansson B. 2017. Sex chromosome evolution: historical insights and future perspectives. Proc Biol Sci. 284: pii:20162806.

Amaral PP, Neyt C, Wilkins SJ, Askarian-Amiri ME, Sunkin SM, Perkins AC, Mattick JS. 2009. Complex architecture and regulated expression of the *Sox2ot* locus during vertebrate development. RNA 15:2013–2027.

Aulchenko YS, Ripke S, Isaacs A, van Duijn CM. 2007. GenABEL: an R library for genome-wide association analysis. Bioinformatics 23:1294–1296.

Bao L, Tian C, Liu S, Zhang Y, Elaswad A, Yuan Z, Khalil K, Sun F, Yang Y, Zhou T, et al. 2019. The Y chromosome sequence of the channel catfish suggests novel sex determination mechanisms in teleost fish. BMC Biol. 17:6.

Baroiller JF, D’Cotta H. 2019. Sex control in Tilapias. In: Wang HP, Piferrer F, Chen SL, Shen ZG, editors. Sex Control in Aquaculture, vol. II, John Wiley & Sons Ltd. p. 191–234.

Bolger AM, Lohse M, Usadel B. 2004. Trimmomatic: a flexible trimmer for Illumina sequence data. Bioinformatics 30:2114–2120.

Bouza C, Sánchez L, Martínez P. 1994. Karyotypic characterization of turbot (*Scophthalmus maximus*) with conventional, fluorochrome, and restriction endonuclease banding techniques. Mar Biol, 120:609–613.

Bouza C, Hermida M, Pardo BG, Fernández C, Fortes G, Castro J, Sánchez L, Presa P, Pérez M, Sanjuán A, et al. 2007. A microsatellite genetic map of the turbot (*Scophthalmus maximus*). Genetics 177:2457–2467.

Bray NL, Pimentel H, Melsted P, Pachter L. 2016. Near-optimal probabilistic RNA-seq quantification. Nat Biotechnol. 34:525–527.

Bull JJ. 1983. Evolution of Sex Determining Mechanisms. Menlo Park, CA, USA. Benjamin Cummings Press.

Capel B. 2017. Vertebrate sex determination: evolutionary plasticity of a fundamental switch. Nat Rev Genet. 18:675–689.

Charlesworth B. 1996. The evolution of chromosomal sex determination and dosage compensation. Curr Biol. 6:149–162.

Charlesworth D. 2019. Young sex chromosomes in plants and animals. New Phytol (doi: 10.1111/nph.16002).

Charlesworth B, Sniegowski P, Stephan W. 1994. The evolutionary dynamics of repetitive DNA in eukaryotes. Nature 371:215– 220.

Chen WM, Abecasis GR. 2007. Family-based association tests for genomewide association scans. Am J Hum Genet. 81:913–926.

Cioffi MB, Yano CF, Sember A, Bertollo LAC. 2017. Chromosomal evolution in lower vertebrates: sex chromosomes in neotropical fishes. Genes 8:258.

Cruz VP, Vera M, Pardo BG, Taggart J, Martínez P, Oliveira C, Foresti F. 2017. Identification and validation of single nucleotide polymorphisms as tools to detect hybridization and population structure in freshwater stingrays. Mol Ecol Resour. 17:550–556.

Cuñado N, Terrones J, Sánchez L, Martínez P, Santos JL. 2002. Sex-dependent synaptic behaviour in triploid turbot, *Scophthalmus maximus* (Pisces, Scophthalmidae). Heredity 89:460–464.

Devlin RH, Nagahama Y. 2002. Sex determination and sex differentiation in fish: an overview of genetic, physiological, and environmental influences. Aquaculture 208:191–364.

Dobin A, Davis CA, Schlesinger F, Drenkow J, Zaleski C, Jha S, Batut P, Chaisson M, Gingeras TR. 2013. STAR: ultrafast universal RNA-seq aligner. Bioinformatics 29:15–21.

Duggal P, Gillanders EM, Holmes TN, Bailey-Wilson JE. 2008. Establishing an adjusted p-value threshold to control the family-wide type 1 error in genome wide association studies. BMC Genomics 9:516.

Figueras A, Robledo D, Corvelo A, Hermida M, Pereiro P, Rubiolo JA, Gómez-Garrido J, Carreté L, Bello X, Gut M, et al. 2016. Whole genome sequencing of turbot (*Scophthalmus maximus*; Pleuronectiformes): a fish adapted to demersal life. DNA Res. 23:181–192.

Gammerdinger WJ, Kocher TD. 2018. Unusual diversity of sex chromosomes in African cichlid fishes. Genes 9:pii: E480.

Guo L, Xu YH, Zhang N, Zhou FL, Huang JH, Liu BS, Jiang SG, Zhang DC. 2019. A high-density genetic linkage map and QTL mapping for sex in black tiger shrimp (*Penaeus monodon*). Front Genet. 10:326.

Gou Y, Vemaraju S, Sweet EM, Kwon HJ, Riley BB. 2018. *sox2* and *sox3* Play unique roles in development of hair cells and neurons in the zebrafish inner ear. Dev Biol. 435:73–83.

Haffray P, Lebègue E, Jeu S, Guennoc M, Guiguen Y, Baroiller JF, Fostier A. 2009. Genetic determination and temperature effects on turbot *Scophthalmus maximus* sex differentiation: an investigation using steroid sex-inverted males and females. Aquaculture 294:30–36.

Harts AMF, Schwanz LE, Kokko H. 2014. Demography can favour female-advantageous alleles. Proc Biol Sci. 281:20140005.

Hattori RS, Murai Y, Oura M, Masuda S, Majhi SK, Sakamoto T, Fernandino JI, Somoza GM, Yokota M, Strüssmann CA. 2012. A Y-linked anti-Müllerian hormone duplication takes over a critical role in sex determination. Proc Natl Acad Sci USA 109:2955–2959.

Hermida M, Bouza C, Fernández C, Sciara AA, Rodríguez-Ramilo ST, Fernández J, Martínez P. 2013. Compilation of mapping resources in turbot (*Scophthalmus maximus*): A new integrated consensus genetic map. Aquaculture 414–415:19-25.

Kamiya T, Kai W, Tasumi S, Oka A, Matsunaga T, Mizuno N, Fujita M, Suetake H, Suzuki S, Hosoya S, et al. 2012. A Trans-species missense SNP in Amhr2 is associated with sex determination in the tiger pufferfish, *Takifugu rubripes* (fugu). PLoS Genet. 8:e1002798.

Kato Y, Perez CAG, Mohamad Ishak NS, Nong QD, Sudo Y, Matsuura T, Wada T, Watanabe H. 2018. A 5’ UTR-overlapping lncRNA activates the male-determining gene doublesex1 in the crustacean *Daphnia magna*. Curr Biol. 28:1811–1817.

Koyama T, Nakamoto M, Morishima K, Yamashita R, Yamashita T, Sasaki K, Kuruma Y, Mizuno N, Suzuki M, Okada Y, et al. 2019. A SNP in a steroidogenic enzyme is associated with phenotypic sex in *Seriola* fishes. Curr Biol. 29:1901–1909.

Koren S, Walenz BP, Berlin K, Miller JR, Bergman NH, Phillippy AM. 2017. Canu: scalable and accurate long-read assembly via adaptive k-mer weighting and repeat separation. Genome Res. 27:722–736.

Lasne C, Sgrò CM, Connallon T. 2017. The relative contributions of the X chromosome and autosomes to local adaptation. Genetics 205:1285–1304.

Li H. 2013. Aligning sequence reads, clone sequences and assembly contigs with BWA-MEM. *arXiv*:1303.3997 [q-bio.GN].

Li H, Handsaker B, Wysoker A, Fennell T, Ruan J, Homer N, Marth G, Abecasis G, Durbin R, 1000 Genome Project Data Processing Subgroup. 2009. The Sequence Alignment/Map format and SAMtools. Bioinformatics 25:2078–2079.

Liang S, Liu D, Li X, Wei M, Yu X, Li Q, Ma H, Zhang Z, Qin Z. 2019. *SOX2* participates in spermatogenesis of Zhikong scallop *Chlamys farreri*. Sci Rep. 9:76.

Lima L, Marchet C, Caboche S., Da Silva C, Istace B, Aury JM, Touzet H, Chikhi R. 2019. Comparative assessment of long-read error correction software applied to Nanopore RNA-sequencing data. Brief Bioinform, bbz058, https://doi.org/10.1093/bib/bbz058

Lozano CA, Gjerde B, Odegard J, Bentsen HB. 2013. Heritability estimates for male proportion in the GIFT Nile tilapia (*Oreochromis niloticus* L.). Aquaculture 372:137–148.

Maroso F, Hermida M, Millán A, Blanco A, Saura M, Fernández A, Dalla Rovere M, Bargelloni L, Cabaleiro S, Villanueva B, et al. 2018. Highly dense linkage maps from 31 full-sibling families of turbot (*Scophthalmus maximus*) provide insights into recombination patterns and chromosome rearrangements throughout a newly refined genome assembly. DNA Res. 25:439–450.

Martínez P, Bouza C, Hermida M, Fernández J, Toro MA, Vera M, Pardo BG, Millán A, Fernández C, Vilas R, et al. 2009. Identification of the major sex-determining region of turbot (*Scophthalmus maximus*). Genetics 183:1443–1452.

Martínez P, Viñas AM, Sánchez L, Díaz N, Ribas L, Piferrer F. 2014. Genetic architecture of sex determination in fish: Applications to sex ratio control in aquaculture. Front Genet. 5:340.

Martínez P, Robledo D, Rodríguez-Ramilo ST, Hermida M, Taboada X, Pereiro P, Rubiolo JA, Ribas L, Gómez-Tato A, Alvarez-Dios JA, et al. 2016. Turbot (*Scophthalmus maximus*) genomic resources: Application for boosting aquaculture production. In: MacKenzie S, Jentoft S, editors. Genomics in Aquaculture. London: Elsevier. p. 131–163.

Matsuda M, Nagahama Y, Shinomiya A, Sato T, Matsuda C, Kobayashi T, Morrey CE, Shibata N, Asakawa S, Shimizu N, et al. 2002. DMY is a Y-specific DM-domain gene required for male development in the medaka fish. Nature 417:559–563.

Messemaker TC, van Leeuwen SM, van den Berg PR, ’t Jong AEJ, Palstra RJ, Hoeben RC, Semray S, Mikkers HMM. 2018. Allele-specific repression of *Sox2* through the long non-coding RNA *Sox2ot*. Sci Rep. 8:386.

Matsuda M, Sakaizumi M. 2016. Evolution of the sex-determining gene in the teleostean genus Oryzias. Gen Comp Endocrinol. 239:80–88.

Mulvey BB, Olcese U, Cabrera JR, Horabin JI. 2014. An interactive network of long non-coding RNAs facilitates the *Drosophila* sex determination decision. Biochim Biophys Acta. 1839:773–784.

Myosho T, Otake H, Masuyama H, Kuroki Y, Fujiyama A, Naruse K, Hamaguchi S, Sakaizumi M. 2012. Tracing the emergence of a novel sex-determining gene in medaka, *Oryzias luzonensis*. Genetics 191:163–170.

Nagabhushana A, Mishra RK. 2016. Finding clues to the riddle of sex determination in zebrafish. J Biosci. 41:145–155.

Ospina-Alvarez N, Piferrer F. 2008. Temperature-dependent sex determination in fish revisited: prevalence, a single sex ratio response pattern, and possible effects of climate change. PLoS One 3:e2837.

Palaiokostas C, Bekaert M, Khan MGQ, Taggart JB, Gharbi K, McAndrew BJ, Penman DJ. 2013. Mapping and validation of the major sex-determining region in Nile Tilapia (*Oreochromis niloticus* L.) using RAD sequencing. PLoS One 8:e68389.

Pan Q, Feron R, Yano A, Guyomard R, Jouanno E, Vigouroux E, Wen M, Busnel JM, Bobe J, Concordet JP, et al. 2019. Identification of the master sex determining gene in Northern pike (*Esox lucius*) reveals restricted sex chromosome differentiation. PLoS Genet. 15: e1008013.

Parnell NF, Streelman JT. 2013. Genetic interactions controlling sex and color establish the potential for sexual conflict in Lake Malawi cichlid fishes. Heredity 110: 239–246.

Peichel CL, Ross JA, Matson CK, Dickson M, Grimwood J, Schmutz J, Myers RM, Mori S, Schluter D, Kingsley DM. 2004. The master sex-determination locus in threespine sticklebacks is on a nascent Y chromosome. Curr Biol. 14:1416–1424.

Penman DJ, Piferrer F. 2008. Fish gonadogenesis. Part I. Genetic and environmental mechanisms of sex determination. Rev Fish Sci. 16(S1):16–34.

Perrin N. 2016. Random sex determination: When developmental noise tips the sex balance. Bioessays 38:1218–1226.

Piferrer F, Cal RM, Gómez C, Alvarez-Blázquez B, Castro J, Martínez P. 2004. Induction of gynogenesis in the turbot (*Scophthalmus maximus*): Effects of UV irradiation on sperm motility, the Hertwig effect and viability during the first 6 months of age. Aquaculture 238: 403–419.

Piferrer F, Martínez P, Ribas L, Viñas A, Díaz N. 2012. Functional genomic analysis of sex determination and differentiation in teleost fish. In: Saroglia M, Liu Z, editors. Functional Genomics in Aquaculture. Oxford:Wiley-Blackwell. p. 169–204.

Ribas L, Robledo D, Gómez-Tato A, Viñas A, Martínez P, Piferrer F. 2016. Comprehensive transcriptomic analysis of the process of gonadal sex differentiation in the turbot (*Scophthalmus maximus*). Mol Cell Endocrinol. 422:132–149.

Robinson JT, Thorvaldsdóttir H, Winckler W, Guttman M, Lander ES, Getz G, Mesirov JP. 2011. Integrative genomics viewer. Nat Biotechnol. 29:24–26.

Robledo D, Hernández-Urcera J, Cal RM, Pardo BG, Sánchez L, Martínez P, Viñas A. 2014. Analysis of qPCR reference gene stability determination methods and a practical approach for efficiency calculation on a turbot (*Scophthalmus maximus*) gonad dataset. BMC Genomics 15:648.

Robledo D, Ribas L, Cal R, Sánchez L, Piferrer F, Martínez P, Viñas A. 2015. Gene expression analysis at the onset of sex differentiation in turbot (*Scophthalmus maximus*). BMC Genomics 16:973.

Robledo D, Palaiokostas C, Bargelloni L, Martínez P, Houston R. 2018. Applications of genotyping by sequencing in aquaculture breeding and genetics. Rev Aquacult. 10:670–682.

Rousset F. 2008. genepop’007: a complete re-implementation of the genepop software for Windows and Linux. Mol Ecol Resour. 8:103–106.

Shahryari A, Jazi MS, Samaei NM, Mowla SJ. 2015. Long non-coding RNA SOX2OT: expression signature, splicing patterns, and emerging roles in pluripotency and tumorigenesis. Front Genet. 6:196.

Ser JR, Roberts RB, Kocher TD. 2010. Multiple interacting loci control sex determination in lake Malawi cichlid fish. Evolution 64:486–501.

Star B, Tørresen OK, Nederbragt AJ, Jakobsen KS, Pampoulie C, Jentoft S. 2016. Genomic characterization of the Atlantic cod sex-locus. Sci Rep. 6:31235.

Steevens AR, Glatzer JC, Kellogg CC, Low WC, Santi PA, Kiernan AE. 2019. SOX2 is required for inner ear growth and cochlear nonsensory formation before sensory development. Development 146: dev170522.

Taboada X, Pansonato-Alves JC, Foresti F, Martínez P, Viñas A, Pardo BG, Bouza C. 2014. Consolidation of the genetic and cytogenetic maps of turbot (*Scophthalmus maximus*) using FISH with BAC clones. Chromosoma 123: 281–291.

Taboada X, Robledo D, Bouza C, Piferrer F, Viñas AM, Martínez P. 2019. Reproduction and sex control in turbot. In: Wang HP, Piferrer F, Chen SL, Shen ZG, editors. Sex Control in Aquaculture, vol. II, John Wiley & Sons Ltd. p. 565–582.

Tan MH, Austin CM, Hammer MP, Lee YP, Croft LJ, Gan HM. 2018. Finding Nemo: hybrid assembly with Oxford Nanopore and Illumina reads greatly improves the clownfish (*Amphiprion ocellaris*) genome assembly. Gigascience 7:1–6.

Takehana Y, Matsuda M, Myosho T, Suster ML, Kawakami K, Shin-I T, Kohara Y, Kuroki Y, Toyoda A, Fujiyama A, et al. 2014. Co-option of Sox3 as the male-determining factor on the Y chromosome in the fish *Oryzias dancena*. Nat Commun. 5:4157.

Tomaszkiewicz M, Medvedev P, Makova KD. 2017. Y and W chromosome assemblies: approaches and discoveries. Trends Genet. 33: 266–282.

Trapnell C, Williams BA, Pertea G, Mortazavi A, Kwan G, van Baren MJ, Salzberg SL, Wold BJ, Pachter L. 2010. Transcript assembly and quantification by RNA-Seq reveals unannotated transcripts and isoform switching during cell differentiation. Nat Biotechnol. 28:511–515.

Utsunomia R, Scacchetti PC, Hermida M, Fernández-Cebrián R, Taboada X, Fernández C, Bekaert M, Mendes NJ, Robledo D, Mank JE, et al. 2017. Evolution and conservation of *Characidium* sex chromosomes. Heredity 119:237–244.

Vandeputte M, Dupont-Nivet M, Chavanne H, Chatain B. 2007. A polygenic hypothesis for sex determination in the European sea bass *Dicentrarchus labrax*. Genetics 176:1049–1057.

Vaser R, Sović I, Nagarajan N, Šikić M. 2017. Fast and accurate de novo genome assembly from long uncorrected reads. Genome Res. 27:737–746.

Viñas A, Taboada X, Vale L, Robledo D, Hermida M, Vera M, Martínez P. 2012. Mapping of DNA sex-specific markers and genes related to sex differentiation in turbot (*Scophthalmus maximus*). Mar Biotechnol. 14:655–663.

Walker BJ, Abeel T, Shea T, Priest M, Abouelliel A, Sakthikumar S, Cuomo CA, Zeng Q, Wortman J, Young SK, et al. 2014. Pilon: an integrated tool for comprehensive microbial variant detection and genome assembly improvement. PLoS One 9:e112963.

Wang S, Meyer E, McKay JK, Matz MV. 2012. 2b-RAD: a simple and flexible method for genome-wide genotyping. Nat Methods 9:808–810.

Xing Y, Liu Y, Zhang Q, Nie X, Sun Y, Zhang Z, Li H, Fang K, Wang G, Huang H, et al. 2019. Hybrid *de novo* genome assembly of Chinese chestnut (*Castanea mollissima*), GigaScience 8:giz112.

Yano A, Nicol B, Jouanno E, Quillet E, Fostier A, Guyomard R, Guiguen Y. 2013. The sexually dimorphic on the Y-chromosome gene (sdY) is a conserved male-specific Y-chromosome sequence in many salmonids. Evol Appl. 6:486–496.

Yano CF, Bertollo LA, Ezaz T, Trifonov V, Sember A, Liehr T, Cioffi M.B. 2016. Highly conserved Z and molecularly diverged W chromosomes in the fish genus Triportheus (Characiformes, Triportheidae). Heredity 118: 276–283.

Zhou HY, Katsman Y, Dhaliwal NK, Davidson S, Macpherson NN, Sakthidevi M, Collura F, Mitchell JA. 2014. A *Sox2* distal enhancer cluster regulates embryonic stem cell differentiation potential. Genes Dev. 28:2699–2711.

